# SpxA1 and SpxA2 Function as a Stoichiometry-Dependent Regulatory Rheostat Governing Virulence Gene Expression in Group A Streptococcus

**DOI:** 10.64898/2026.06.19.733365

**Authors:** Gretchen A. Morrison, Luis Alberto Vega, Misú Sanson Iglesias, Khufu Holly, W. Hayes McDonald, Nicola Horstmann, Samuel A. Shelburne, Jennifer A. Gaddy, Anthony R. Flores

## Abstract

Group A Streptococcus (GAS) is a human-restricted pathogen whose global incidence has surged in the post-COVID era. The ability of GAS to shift from a colonizing to invasive phenotype depends on coordinated virulence gene regulation in response to host-derived signals. However, the mechanisms by which individual stress-sensing systems interact to reshape the virulence gene regulatory landscape remain incompletely understood. Here we define the regulatory programs of two conserved transcriptional regulator paralogs, SpxA1 and SpxA2, using an integrated multi-omic approach combining RNA-seq, data-independent acquisition proteomics, NanoString-based transcriptional profiling across multiple host-relevant stress conditions, and chromatin immunoprecipitation with exonuclease treatment (ChIP-exo). RNA-seq revealed functionally distinct regulons with SpxA1 governing oxidative stress defense and SpxA2 coordinating virulence-associated gene expression linked to the CovRS two-component regulatory system. Proteomic analysis established SpxA2 as a ClpXP protease substrate in GAS and identified reciprocal paralog accumulation upon loss of either SpxA1 or SpxA2, consistent with compensatory transcriptional upregulation. NanoString profiling under bacitracin and human neutrophil peptide-1 challenge identified four gene modules with distinct stoichiometry-dependent and condition-dependent regulatory logic, revealing that the SpxA1/SpxA2 ratio rather than the activity of either paralog alone determines which transcriptional programs are engaged. ChIP-exo demonstrated that SpxA2 directly modulates CovR-DNA binding occupancy in a CovR binding motif-dependent manner, simultaneously antagonizing CovR dimer binding at an extended (25-bp) CovR motif and facilitating CovR monomer binding at the canonical ATTARA motif. These findings establish the LiaFSR-SpxA2-CovRS axis as a cross-regulatory circuit through which GAS cell envelope stress sensing is directly transduced into coordinated virulence gene regulatory changes.

**Importance:** Group A *Streptococcus* (GAS) causes millions of infections annually, including a recent global surge in invasive disease. To survive in the human host, GAS must rapidly reprogram virulence gene expression in response to host-derived stresses. This study characterizes two conserved transcriptional regulators, SpxA1 and SpxA2, that govern this response through interaction with RNA polymerase to indirectly influence the DNA-binding activity of downstream transcription factors. We show that SpxA2, activated by a cell envelope stress-sensing system responding to human antimicrobial peptides, reshapes the binding of the master virulence regulator CovR in a promoter-specific manner, coupling cell envelope stress sensing to virulence gene regulation. The stoichiometric balance between SpxA1 and SpxA2 functions as a regulatory rheostat calibrating overall virulence gene regulatory tone, providing a framework for understanding how RNA polymerase-interacting regulators coordinate stress responses and virulence gene control across Gram-positive bacterial pathogens.

## Introduction

Group A *Streptococcus* (GAS, *Streptococcus pyogenes*) is a human-specific pathogen responsible for more than 700 million infections worldwide annually (1). GAS causes a broad spectrum of disease, from relatively benign superficial infections such as pharyngitis to severe, life-threatening invasive disease including necrotizing fasciitis and streptococcal toxic shock syndrome, which carry case fatality rates of 10–20% (1, 2). Invasive GAS disease (iGAS) infections more than doubled between 2013 and 2022 in the US (3) and several countries reported marked surges in invasive GAS disease incidence during the later stages of the COVID-19 pandemic (4–7) highlighting the urgent need to better define the mechanisms by which this pathogen causes disease. Despite its clinical importance, the molecular mechanisms governing GAS virulence gene regulation in response to host-derived signals remain incompletely understood.

GAS encodes a complex regulatory network comprising 13 two-component systems (TCSs) and at least 30 stand-alone transcriptional regulators that collectively govern virulence gene expression (8). The LiaFSR (lipid II-interacting antibiotic) three-component system (3CS), conserved across Gram-positive bacteria, responds to cell envelope stress (CES) induced by antimicrobial peptides (AMPs) and cell wall-active antibiotics (9, 10). In GAS, LiaFSR is specifically activated by the α-defensin human neutrophil peptide-1 (hNP-1) through Lipid II-dependent disruption of the cell envelope (11, 12) leading to LiaR phosphorylation and direct transcriptional activation of *spxA2* (12, 13). LiaR-dependent *spxA2* induction is conserved in *Streptococcus mutans* (14, 15), establishing this regulatory connection as a conserved feature of streptococcal CES responses. The restricted LiaR regulon in GAS implies a greater role of SpxA2 in downstream gene regulation but remains undefined.

The control of virulence regulatory system (CovRS, also known as CsrRS) is the most extensively characterized TCS in GAS and functions as a master regulator of virulence gene expression. CovS functions as both a kinase and a phosphatase for CovR, with the characteristics of the external signal determining the balance of activity. Mg^2+^ promotes CovR phosphorylation and repression of virulence genes (16), while LL-37 enhances CovS phosphatase activity to de-repress CovR-regulated virulence gene expression (17) – a pattern inverted relative to canonical TCS activation. Recent ChIP-seq in the *emm3* strain MGAS10870 defined two distinct patterns of CovR-DNA occupancy: an extended (25-bp) AT-rich pseudopalindromic consensus sequence (WTWTTATAAWAAAAWNATDA) consistent with dimeric CovR binding, and a shorter canonical ATTARA motif suggestive of monomeric CovR interaction (18). Prior work from our laboratory demonstrated that *spxA2* transcript levels correlate with CovRS-regulated virulence gene expression (12), suggesting a regulatory interaction between the LiaFSR-SpxA2 axis and CovRS-dependent virulence gene control.

The Spx (*<u>s</u>*uppressor of Clp*<u>P</u>* and Clp*<u>X</u>*) proteins are a family of transcriptional regulators highly conserved across the Bacillota. First characterized in *Bacillus subtilis*, Spx binds to the C-terminal domain of the RNAP α subunit (αCTD) and modulates transcription of stress-responsive gene sets (19, 20). Orthologous Spx proteins function in oxidative stress responses, cell envelope homeostasis, and virulence gene regulation across low-GC Gram-positive pathogens (21–29). Many pathogenic streptococci possess two Spx paralogs suggesting distinct regulatory roles (30). In *S. mutans*, SpxA1 governs oxidative stress gene regulation while SpxA2 coordinates cell envelope homeostasis downstream of LiaFSR, with differential ClpXP-mediated proteolysis contributing to paralog-specific abundance dynamics (15, 21, 22). In GAS, Port et al. demonstrated individual roles of SpxA1 and SpxA2 in the GAS stress responses and virulence (31). Akin to that in *S. mutans* (14), we showed that the GAS LiaR directly activates *spxA2* transcription but in specific response to hNP-1-mediated CES (12, 13). Despite the extensive work in CovRS and Spx paralogs in GAS and related organisms, the mechanism by which SpxA2 influences CovR-dependent gene expression and the extent to which the paralogs act cooperatively or antagonistically across physiologically relevant stress conditions are not fully resolved.

The present study defines the transcriptional regulatory roles of SpxA1 and SpxA2 in GAS using a multi-omics approach. SpxA1 and SpxA2 have functionally distinct and largely non-overlapping regulons, with SpxA1 governing oxidative stress defense and SpxA2 coordinating a broad virulence gene regulatory program. NanoString profiling identified four gene modules with distinct stoichiometry- and condition-dependent gene regulation. We also report the first use of high-resolution ChIP-exo in GAS to reveal that SpxA2 directly modulates CovR-DNA binding in a motif-dependent manner. The work presented here establishes the molecular basis for LiaFSR-SpxA2-CovRS cross-regulatory control of GAS virulence gene expression and defines the SpxA1/SpxA2 balance as a regulatory rheostat calibrating the overall virulence gene regulatory tone of the GAS cell.

## Results

### Δ*spxA1* and Δ*spxA2* have distinct transcriptomes

SpxA1 and SpxA2 were previously shown to have opposing roles in virulence and the oxidative stress response in GAS (31). To define the regulons of SpxA1 and SpxA2, we performed RNA-seq comparing the parental *emm3* MGAS10870 (WT) to isogenic mutant strains Δ*spxA1* and Δ*spxA2* during exponential-phase growth. Analysis of differential gene expression (DGE; ≥1.5-fold change, *P*-adj ≤ 0.05 by Benjamini-Hochberg correction, core genome excluding prophage and mobile elements) revealed unique mutant transcriptomes (Table 1, Figure 1; RNA-seq quality metrics in Supplemental Figure S1). The Δ*spxA1* mutant showed a substantially smaller regulon (158 genes) compared to Δ*spxA2* (419 genes); the two transcriptomes share 77 differentially expressed genes (53 co-downregulated, 18 co-upregulated, 6 opposing) consistent with both shared and paralog-specific gene regulation (Supplemental Table ST1).

**Figure 1.**
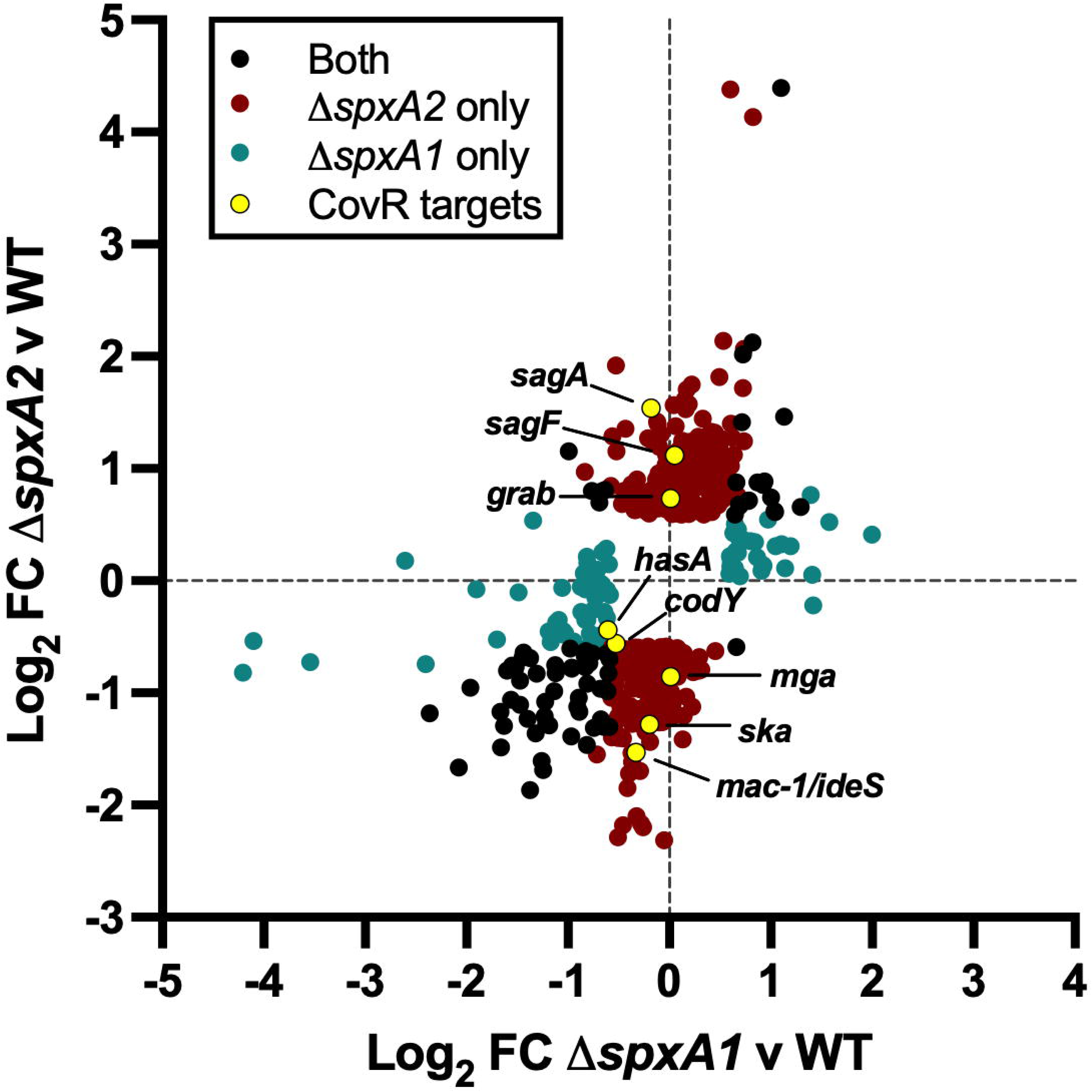
Δ*spxA1* and Δ*spxA2* have distinct transcriptomes. Scatter plot comparing genome-wide differential gene expression (DGE) in Δ*spxA1* (x-axis) and Δ*spxA2* (y-axis) relative to the parental strain MGAS10870 (WT). Each point represents a single gene meeting significance criterion (|log_2_FC| ≥ log_2_(1.5), *P*-adj ≤ 0.05, Benjamini-Hochberg correction) in at least one mutant background. Genes differentially expressed in both mutants are shown in black (n=77); genes differentially expressed exclusively in Δ*spxA2* are shown in red (n=342); genes differentially expressed exclusively in Δ*spxA1* are shown in teal (n=81). Select CovR-regulated targets are highlighted in yellow and genes labeled. Full RNA-seq differential expression results are provided in Supplemental Table ST1.

**Table 1.** Select differently expressed genes unique to either the Δ*spxA1* or Δ*spxA2* isogenic mutants relative to that in the parental strain, MGAS10870.

| **$\Delta spxA1$ vs MGAS10870** |  |  |  |
| --- | --- | --- | --- |
| **Locus tag<sup>a</sup>** | **Gene<sup>b</sup>** | **Predicted product** | **FC<sup>c</sup>** |
| C9Q_01310 (SpyM3_0208) | *sufC* | Fe-S cluster assembly ATPase | -2.25 |
| C9Q_06895 (SpyM3_1363) | *cysK* | Cysteine synthase A | +2.21 |
| C9Q_05400 (SpyM3_1071) | *sodA* | Superoxide dismutase | -3.74 |
| C9Q_04415 (SpyM3_0808) | *nox.1* | Putative NADH oxidase | -2.93 |
| **$\Delta spxA2$ vs MGAS1870** |  |  |  |
| C9Q_08815 (SpyM3_1742) | *speB* | Cysteine protease | +21.1 |
| C9Q_03565 (SpyM3_0583) | ***ideS*** | IgG protease (Mac-1) | -2.89 |
| C9Q_08605 (SpyM3_1698) | ***ska*** | Streptokinase A precursor | -2.43 |
| C9Q_05215 (SpyM3_1032) | ***grab*** | Protein G-related α2-macroglobulin-binding protein | +1.69 |
| C9Q_02050 (SpyM3_0346) | *rgg2* | Quorum-sensing transcriptional regulator | -2.71 |
| C9Q_09025 (SpyM3_1784) | ***pepO*** | Putative endopeptidase O | -2.21 |
| C9Q_03070 (SpyM3_0485) | ***sagF*** | Streptolysin associated protein | +2.17 |
| C9Q_06010 (SpyM3_1196) | *arcA* | Arginine deiminase | -4.95 |

<sup>a</sup> Locus tag designation reflects current MGAS10870 genome annotation (GenBank NZ\_CP067090.1; C9Q prefix). Corresponding prior SpyM3 locus tags shown in parentheses where available.
<sup>b</sup> Gene names in bold font indicate predicted or known regulation by CovRS.
<sup>c</sup> Fold-change (FC) values calculated from DESeq2 $\log_2$ FC (linear scale); $P$ -adj < 0.05 (Benjamini-Hochberg correction). Complete differential expression results are provided in Supplemental Table S1.

The composition of the Δ*spxA1* and Δ*spxA2* transcriptomes was consistent with previously described divergent roles of Spx paralogs (21, 22). In Δ*spxA1*, DGE was enriched for oxidative stress management (*sodA*, *nox.1*) and redox homeostasis (*sufC*, *cysK*) genes (Table 1). The cysteine protease, *speB*, was the most highly upregulated gene in the Δ*spxA2* transcriptome (+21-fold; Table 1), consistent with prior observations (12). Several Δ*spxA2* DGE genes are known to be regulated by the CovRS TCS (Table 1, bold font; Figure 1, yellow dots with labels), including *ideS/mac-1* and *ska*. We previously showed a direct correlation between *spxA2* transcript level and CovR-dependent gene regulation (12), consistent with a regulatory axis linking LiaFSR, SpxA2, and CovRS-dependent virulence gene expression (Table 1, gene names in bold font). In total, the transcriptome data define distinct SpxA1 and SpxA2 regulatory programs: SpxA1 governing oxidative stress defense and SpxA2 governing CovRS cross-regulated virulence gene expression.

### SpxA2 is required for the response to AMP-mediated cell envelope stress

The distinct regulon compositions led us to hypothesize that SpxA1 and SpxA2 respond differently to AMP-mediated cell envelope stress. AMP tolerance was assessed following bacitracin and hNP-1 exposure *in vitro* using virtual colony counts (CFU_v_) (13) of WT, Δ*spxA1*, and Δ*spxA2*. We observed significantly reduced AMP tolerance in Δ*spxA2* but not in Δ*spxA1* (Figure 2A), indicating that SpxA2 alone is required for AMP tolerance *in vitro*.

**Figure 2.**
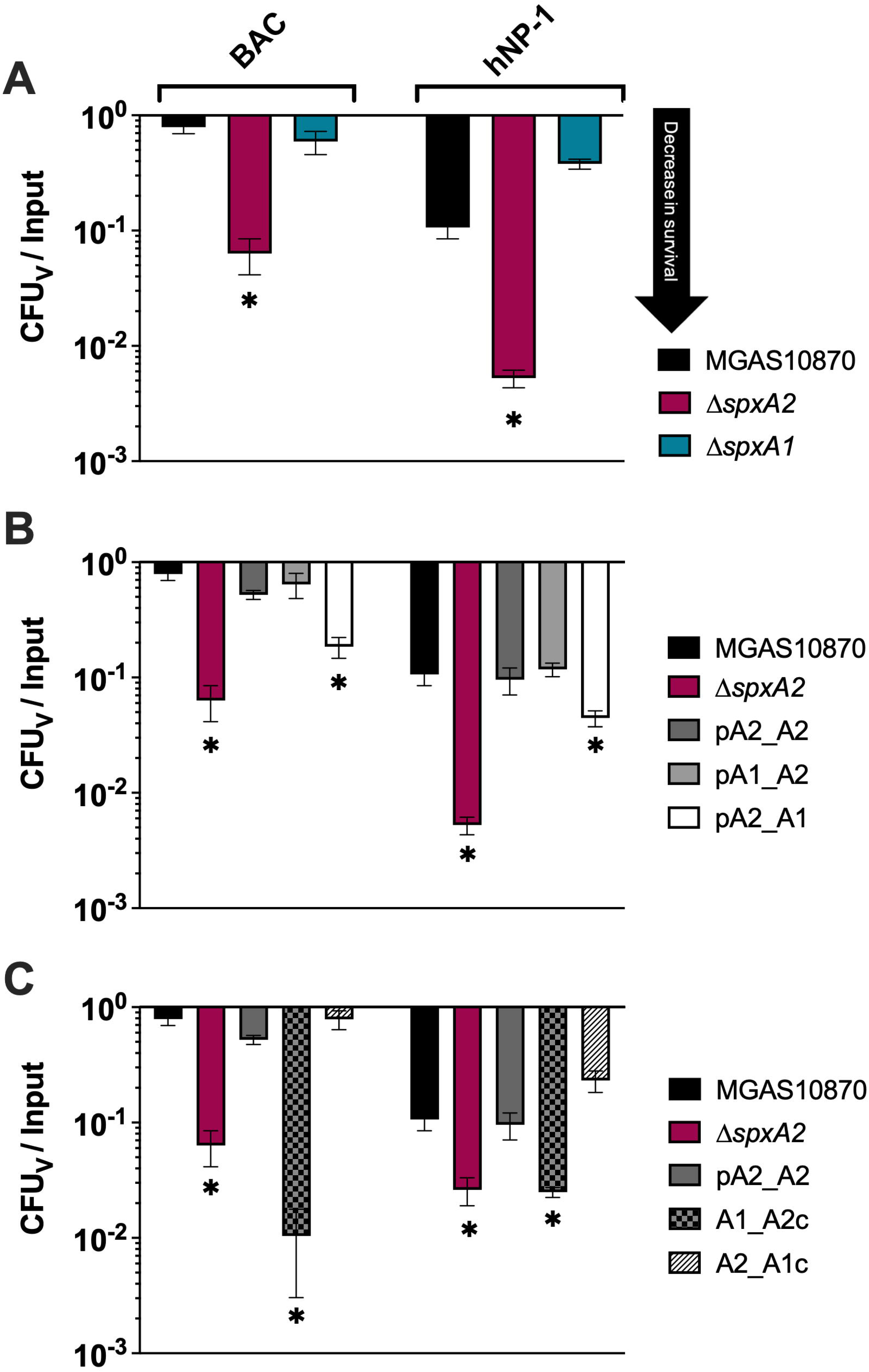
SpxA2 is required for tolerance to AMP-mediated cell envelope stress. Tolerance to bacitracin (BAC, 1 μg/mL) and human neutrophil peptide-1 (hNP-1, 5 μg/mL) was assessed using virtual colony counts (CFU_v_/Input) as described in Methods. AMP stress labeled at the top of panel A are also extended to panels B and C. (**A**) Comparison of AMP tolerance in WT (MGAS10870, black), Δ*spxA2* (red), and Δ*spxA1* (teal). Significantly reduced tolerance only observed in Δ*spxA2*. (**B**) AMP tolerance in Δ*spxA2* expressing promoter-swapped *spxA1* or *spxA2 in trans*. Expressing *spxA2* from the *spxA2* promoter (pA2_A2, dark gray) or the *spxA1* promoter (pA1_A2, medium gray) restores WT-level tolerance. Expressing spxA1 from the spxA2 promoter (pA2_A1, white) does not restore tolerance. (**C**) AMP tolerance in Δ*spxA2* expressing C-terminal swapped constructs *in trans*. Tolerance is restored by the full SpxA2 complement (pA2_A2, dark gray) and SpxA2 carrying the SpxA1 C-terminus (A2_A1c, hatched), but not by SpxA1 carrying the SpxA2 C-terminus (A1_A2c, checkered). For all panels, bars closer to the x-axis indicate decreased survival (arrow, panel A). Data are presented as mean ± 95% confidence interval from biological triplicate assays. Asterisks indicate significant difference from MGAS10870 (*P* < 0.05, Mann-Whitney U test).

Given that AMP stress results in direct LiaR regulation of *spxA2* transcription via binding to the *spxA2* promoter (12, 13) and not that of *spxA1*, we hypothesized that increased SpxA1 levels would be insufficient to restore AMP tolerance in the Δ*spxA2* background. To test this, we created promoter-swapped strains expressing *spxA1* or *spxA2* from the noncognate promoter *in trans* in the Δ*spxA2* background. Expressing SpxA1 from the *spxA2* promoter (pA2_A1; induction under AMP stress) did not restore AMP tolerance to WT levels, while expressing SpxA2 from the *spxA1* promoter (pA1_A2; no induction under AMP stress) fully restored tolerance (Figure 2B), demonstrating that even low-level constitutive SpxA2 expression complements the CES response and that the divergence between paralogs extends beyond promoter-driven expression differences.

Differential ClpXP-mediated degradation is defined by the C-terminal sequence in SpxA1 and SpxA2 in *S. mutans* and contributes to paralog-specific phenotypes *in vitro* (15). We hypothesized that C-terminal sequence differences may similarly contribute to distinct AMP tolerance phenotypes in GAS. Thus, we next generated C-terminal swapped strains *in trans* expressed from their cognate promoter in the Δ*spxA2* background: SpxA1 with the SpxA2 C-terminus (A1_A2c) and SpxA2 with the SpxA1 C-terminus (A2_A1c) (Supplemental Figure S2). AMP tolerance was restored only by the full SpxA2 complement (pA2_A2) and A2_A1c, but not A1_A2c (Figure 2C). Collectively, these data establish that the SpxA2 functional domain is both necessary and sufficient for AMP tolerance, independent of C-terminal sequence identity.

### Spx protein levels correlate with mutant background

Given the observed complementation phenotypes (Figure 2) and the known susceptibility of Spx proteins to ClpXP-mediated degradation across Gram-positive species (15, 19, 27), we assessed SpxA1 and SpxA2 protein levels in WT, Δ*spxA1*, Δ*spxA2*, Δ*clpX*, and the *in trans* complemented strains by data independent acquisition (DIA) proteomics in the absence of CES (See Methods; Figure 3A). SpxA2 was detected at significantly higher levels in both Δ*clpX* and Δ*spxA1* compared to WT (Figure 3B), representing the first demonstration that SpxA2 is a ClpXP substrate in GAS and extending prior observations from *B. subtilis*, *S. aureus*, and *S. mutans* (15, 19, 27). SpxA1 also trended higher in Δ*clpX* (not significant; Figure 3C), with differential ClpXP efficiency between the paralogs consistent with their C-terminal sequence differences (Supplemental Figure S2) (15). SpxA1 trended toward elevation in Δ*spxA2* (Figure 3C), and the parallel pattern of reciprocal accumulation suggests that loss of either paralog triggers compensatory transcriptional upregulation of the remaining paralog. SpxA1 was significantly elevated in the pA2_A1 strain (Figure 3C) confirming robust *in trans* expression. The failure of this strain to rescue Δ*spxA2* AMP tolerance despite high SpxA1 levels further confirms that functional domain differences underlie paralog divergence.

**Figure 3.**
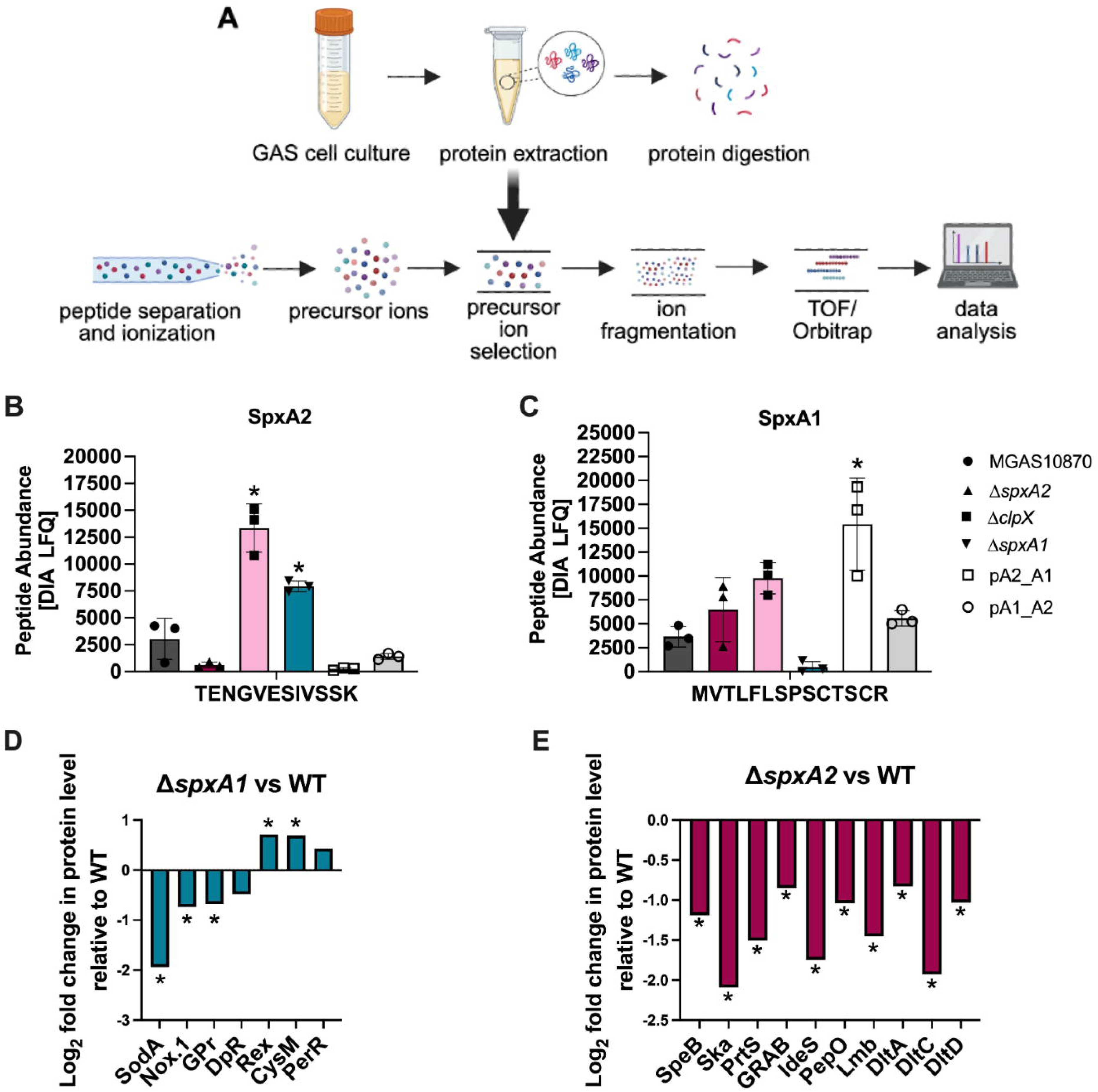
SpxA1 and SpxA2 have distinct proteomic profiles. (**A**) Schematic of DIA proteomics workflow. GAS cell lysates were subjected to tryptic digestion, peptide separation and ionization, precursor ion selection, fragmentation, and mass spectrometric detection for label-free quantification (LFQ). (**B**, **C**) Peptide abundance (DIA LFQ) for representative SpxA2 (TENGVESIVSSK; panel B) and SpxA1 (MVTLFLSPSCTSCR; panel C) tryptic peptides across six strains: MGAS10870 (circle), Δ*spxA2* (triangle), Δ*clpX* (filled square), Δ*spxA1* (inverted triangle), pA2_A1 (open square), and pA1_A2 (open circle). Asterisks indicate significant difference from MGAS10870 (*P* < 0.05, two-tailed ANOVA with Tukey’s post hoc correction). (**D**) Log_2_ fold-change in protein abundance for select oxidative stress defense and redox regulatory proteins in Δ*spxA1* vs. WT. Asterisks indicate significance (*P* < 0.05, limma empirical Bayes moderated *t*-statistic, Benjamini-Hochberg corrected). (**E**) Log_2_ fold-change in protein abundance for select immune evasion and virulence-associated proteins in Δ*spxA2* vs. WT. Asterisks indicate significant differential abundance (*P* < 0.05, limma empirical Bayes moderated *t*-statistic, Benjamini-Hochberg corrected). Complete differential abundance results are provided in Supplemental Table ST2.

### SpxA1 and SpxA2 have distinct proteomic profiles

To define the broader protein-level consequences of SpxA1 and SpxA2 loss, we comprehensively analyzed the DIA proteomics comparing Δ*spxA1* and Δ*spxA2* to the parental MGAS10870. Of 1,269 proteins detected, 218 (17.2%) and 291 (23.4%) showed significant abundance changes in Δ*spxA1* and Δ*spxA2* relative to the WT strain, respectively (|log_2_FC| ≥ log_2_(1.5), *P-*adj < 0.05; Supplemental Table ST2). Proteins with decreased abundance predominated in both mutants: 60% of Δ*spxA1* significant proteins (131/218) and 68% of Δ*spxA2* significant proteins (197/291) were reduced relative to WT (Supplemental Figure S3A-B). Comparison of fold-change values across all 1,269 detected proteins revealed moderate global co-regulation (Pearson r = 0.46; Supplemental Figure S3C), yet the two mutants showed largely non-overlapping sets of significantly changed proteins (87 shared), confirming that SpxA1 and SpxA2 have distinct and non-redundant effects on specific protein targets.

### SpxA1 alters levels of proteins involved in the oxidative stress response

Consistent with the SpxA1 transcriptome and the established oxidative stress regulatory role of SpxA1 in *S. mutans* (22), proteomics revealed significant reductions in SodA and Nox.1 peptide abundance in Δ*spxA1* (Figure 3D) and confirmed concordant transcriptional and translational regulation of these targets. The proteome analysis additionally identified altered abundance of the peroxide resistance proteins GpR and DpR, with GpR significantly reduced and DpR trending towards significance, despite no corresponding transcript-level changes, suggesting SpxA1 more broadly influences the oxidative stress proteome through indirect mechanisms. A candidate indirect mechanism is SpxA1-dependent maintenance of PerR protein levels [the peroxide-sensing master regulator of the GAS oxidative stress response (32)] since the near-significant increase in PerR abundance in Δ*spxA1* (P = 0.053, Figure 3D) may account for downstream proteomic changes without detectable transcript-level effects. Beyond peroxide defense, the significant elevation of Rex (a transcriptional repressor responding to the NAD^⁺^/NADH ratio) in Δ*spxA1* (Figure 3D) indicates a broader shift in redox regulatory homeostasis consistent with loss of SpxA1-dependent Nox.1 activity altering cellular redox balance. Together, the reductions in SodA, Nox.1, DpR, and GPr alongside the trend toward increased PerR and significantly increased Rex suggest that SpxA1 loss produces a broad disruption of redox regulatory homeostasis extending well beyond directly regulated oxidative stress defense genes.

### SpxA2 alters levels of proteins involved in host immune evasion

Proteomics identified significant reductions in peptide abundance for IdeS/Mac-1, Ska, and PepO in Δ*spxA2* (Figure 3E) confirming concordant transcriptional and translational regulation of these immune evasion factors. Additionally, PrtS/SpyCEP, which showed only a trend toward downregulation in RNA-seq, showed a significant reduction at the protein level (Figure 3E), extending SpxA2-dependent regulation of this IL-8 protease to the translational level. The cysteine protease, *speB*, was the most highly upregulated gene in the Δ*spxA2* transcriptome (+21-fold, Table 1) and showed paradoxically decreased peptide abundance, an observation potentially consistent with post-transcriptional regulation of SpeB maturation and secretion (33). Together, these findings support SpxA2-dependent control of CovRS-regulated immune evasion factor protein levels and highlight the complex post-transcriptional regulatory landscape governing individual CovRS targets.

### SpxA1 and SpxA2 differentially regulate GAS transcriptional responses to cell wall and defensin stress

The deletion mutant backgrounds represent fixed extremes of the SpxA1/SpxA2 stoichiometric ratio (Δ*spxA1* is constitutively SpxA2-dominant, while Δ*spxA2* is SpxA1-dominant and cannot be shifted by LiaFSR activation). Thus, we leveraged the isogenic deletion mutants to test how SpxA1/SpxA2 balance shapes the broader GAS transcriptional response to AMP stress. We performed NanoString-based gene expression profiling of WT, Δ*spxA1*, and Δ*spxA2* under uninduced (UI), bacitracin challenge (BAC), and hNP-1 challenge conditions using a validated 116-gene custom codeset (see Methods; Supplemental Table ST3). Differential expression analysis used the limma empirical Bayes framework (34, 35) applied to nSolver-normalized counts (Supplemental Methods; full DGE across all comparisons are in Supplemental Table ST4). One biological replicate (Δ*spxA1* hNP-1, replicate A) was excluded as a global expression outlier by PCA and hierarchical clustering (Supplemental Figures S4-S5) but with minimal impact on model performance given variance estimates informed by 11 additional contrasts (34).

Loss of SpxA2 produced a substantially broader transcriptional response than loss of SpxA1 under both stress conditions (Figure 4A). Under BAC, 53 genes were differentially expressed in Δ*spxA2* compared to 43 in Δ*spxA1* and 38 in WT. Divergence was more pronounced under hNP-1 (46, 23, and 16 genes, respectively). SpxA2 loss was further associated with convergence of the BAC and hNP-1 responses showing 31 genes shared between conditions in Δ*spxA2* compared to 12 and 10 in Δ*spxA1* and WT (Figure 4B). The data indicate that SpxA2 maintains a transcriptional distinction between different types of cell envelope-targeting defensins.

**Figure 4.**
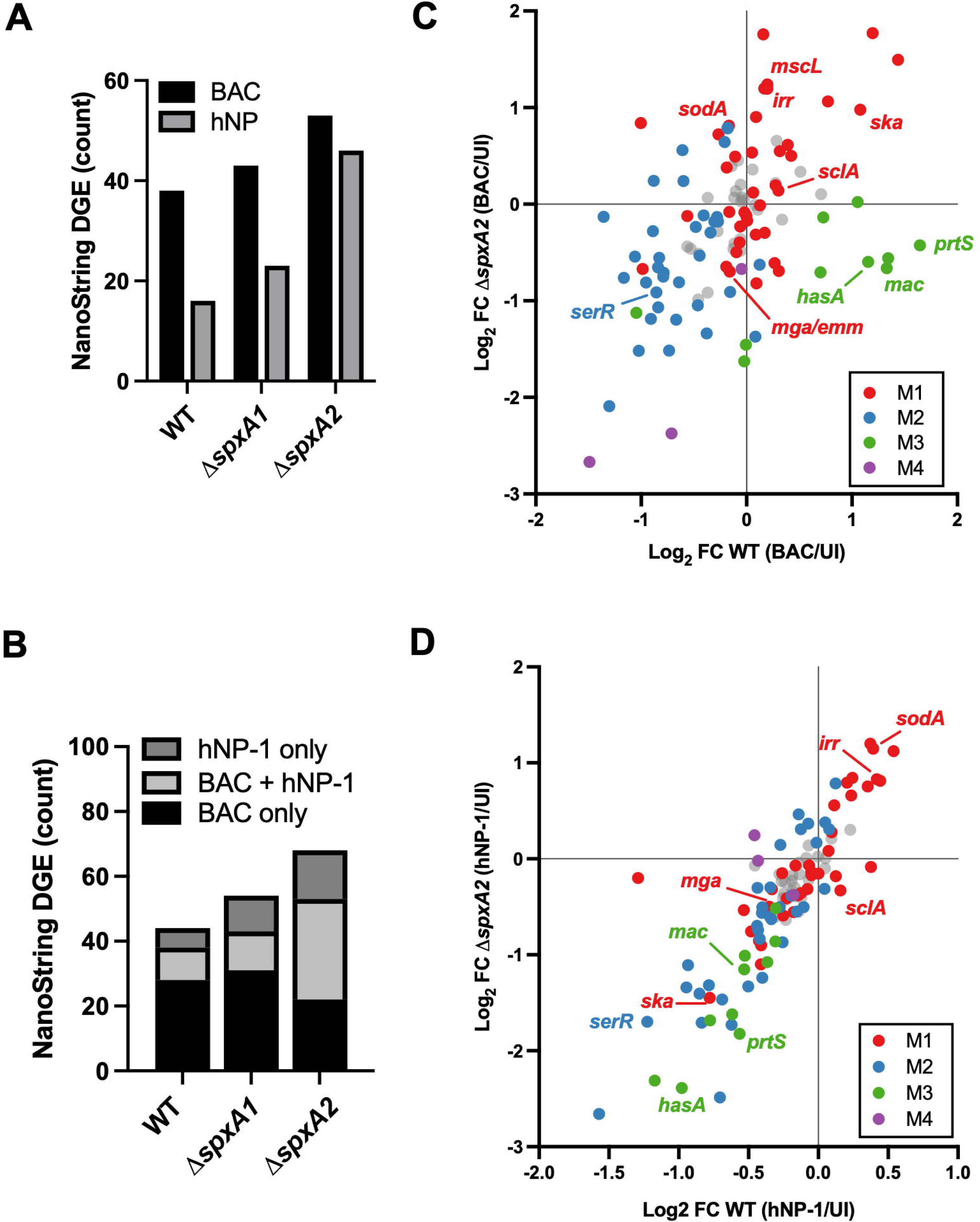
SpxA1 and SpxA2 differentially regulate GAS transcriptional responses to cell wall and defensin stress. NanoString nCounter gene expression profiling was performed in WT (MGAS10870), Δ*spxA1*, and Δ*spxA2* under uninduced (UI), bacitracin (BAC, 1 μg/mL), and hNP-1 (5 μg/mL) conditions using a custom 116-gene codeset (Supplemental Table ST3). Differential expression analysis as described in Results and Supplemental Methods (full results in Supplemental Table ST4). One Δ*spxA1* hNP-1 biological replicate was excluded as a PCA outlier prior to analysis (Supplemental Figures S4-S5). (**A**) Total number of differentially expressed genes (DGE count) under BAC (black) and hNP-1 (gray) for each strain. Δ*spxA2* shows the greatest transcriptional response under both conditions. (**B**) Stacked bar chart showing the proportion of differentially expressed genes that are condition-specific (BAC only, black; hNP-1 only, light gray) or shared between conditions (BAC + hNP-1, dark gray). Δ*spxA2* shows the greatest convergence of BAC and hNP-1 transcriptional responses. (**C**) Scatter plot comparing BAC-induced log_2_FC in WT (x-axis) and Δ*spxA2* (y-axis). Each point represents one NanoString target gene, colored by module assignment: M1 (SpxA1-dependent transcriptional maintenance, red); M2 (SpxA2-dependent repression under non-stress conditions, blue); M3 (Cooperative SpxA1/SpxA2-dependent stress induction, green); M4 (Constitutive SpxA2-dependent repression, purple). Module assignments were derived from unsupervised hierarchical clustering as described in Results and Supplemental Methods (Table 2; Supplemental Table ST5). (**D**) Scatter plot as in panel C comparing hNP-1-induced log_2_FC in WT (x-axis) and Δ*spxA2* (y-axis). Note compressed x-axis range reflecting the more muted WT hNP-1 transcriptional response relative to BAC.

**Table 2.** NanoString nCounter gene expression modules in GAS WT, Δ*spxA1*, and Δ*spxA2* under cell envelope stress conditions.

| Module<sup>a</sup> | Module Name | Features | Genes (n)<sup>b</sup> |
| --- | --- | --- | --- |
| **M1** | SpxA1-dependent transcriptional maintenance | • **Consistently reduced** in Δ*spxA1* vs. WT across all conditions<br>• SpxA1 required for **baseline expression** of diverse gene set<br>• Δ*spxA2* expression broadly unchanged vs. WT | *sodA*, *mscL*, *liaF*, ***codY*** [EM], *ska* [EM], ***mga*** [ATTARA], *dltA* (38) |
| **M2** | SpxA2-dependent repression | • **Elevated** in Δ*spxA2* vs. WT at baseline (UI)<br>• SpxA2 normally **represses** this gene set<br>• BAC/hNP-1 stress **amplifies repression** in both WT and Δ*spxA2* | ***sagA***, ***sagB*** [EM], *nga*, *serR*, *hupY*, *lacC.2*, *manL* (35) |
| **M3** | Cooperative SpxA1/SpxA2-dependent stress induction | • Induced in WT under BAC; **not induced** in either mutant<br>• Both SpxA1 AND SpxA2 required for BAC-induced expression<br>• **Cooperative** rather than paralog-specific regulation | ***hasA*** [EM], ***prtS***, ***scpA*** [EM] *mac*, *arcA* (10) |
| **M4** | Constitutive SpxA2-dependent repression | • **Strongly elevated** in Δ*spxA2* vs. WT regardless of condition<br>• SpxA2 repression is **condition-independent** | *speB*, *SpyM3_0098* (*cbp*), *SpyM3_0100* (*tee3*) (3) |

<sup>a</sup> Module assignments from k=4 hierarchical clustering (Euclidean distance, complete linkage) of log<sub>2</sub>FC profiles across 12 valid NanoString contrasts (See Results and Supplemental Methods for details). Font color corresponds with module designations in Figures 4 and 5.
<sup>b</sup> Complete gene membership with all log<sub>2</sub>FC values provided in Supplemental Table ST5. [EM] = CovR (repressed) target with the extended CovR binding motif (EM); [ATTARA] = CovR (activated) target with the ATTARA-only binding motif, per Horstmann et al. 2023 (18).

We performed unsupervised hierarchical clustering of log_2_FC profiles across all valid contrasts and identified four gene modules (M1-M4) characterized by both SpxA1/SpxA2 imbalance and BAC/hNP-1 stress condition (Table 2; Supplemental Figure S6; Supplemental Table ST5). M1 (SpxA1-dependent transcriptional maintenance; n=38) comprises genes consistently reduced in Δ*spxA1* across all conditions, including oxidative stress defense factors (*sodA*, *mscL*), and regulatory proteins (*liaF*, *codY*, *mga*). M2 (SpxA2-dependent repression; n=35) comprises genes elevated in Δ*spxA2* at baseline and suppressed under stress regardless of SpxA2 status, including *sagA*, *sagB*, *nga*, *serR*, and *hupY*. M3 (Cooperative SpxA1/SpxA2-dependent stress induction; n=10) comprises genes induced in WT under BAC but not induced in either mutant, anchored by CovR-regulated immune evasion genes *hasA*, *prtS*, *scpA*, and *mac-1*. M4 (Constitutive SpxA2-dependent repression; n=3) comprises genes strongly repressed by SpxA2 regardless of condition, anchored by *speB* and two pilus-associated locus proteins (*cbp*, *tee3*). Collectively, the modules are consistent with a model in which the SpxA1:SpxA2 balance dictates GAS regulatory tone and response to stress stimulus.

Comparison of WT and Δ*spxA2* log_2_FC profiles revealed two mechanistically distinct consequences of SpxA2 loss. Under BAC, M3 module genes were induced in WT but failed to reach equivalent induction in Δ*spxA2*, demonstrating that SpxA2 is required for transcriptional activation of CovR-regulated immune evasion genes under cell wall stress (Figure 4C, green dots). Concurrently, M1 module genes including *sodA*, *irr*, and *mscL* showed higher induction in Δ*spxA2* than WT, indicating SpxA2 normally moderates the BAC-induced expression of these targets (Figure 4C, red dots). Under hNP-1, M3 module genes including *hasA*, *prtS*, and *mac-1*, remained more strongly downregulated in Δ*spxA2* than WT (Figure 4D), consistent with SpxA2 being required for the induction of CovR-regulated immune evasion gene expression under both BAC and defensin challenge. Together, the data support a model in which SpxA2 functions as a bidirectional transcriptional regulator under stress, promoting cooperative induction of CovR-regulated immune evasion genes (M3) while moderating a secondary stress response gene set (M1) under both BAC and hNP-1 challenge.

Direct comparison of Δ*spxA1* and Δ*spxA2* log_2_FC profiles provides independent visual validation of the four-module regulatory architecture and its stoichiometric basis (Figure 5A-B). Under BAC stress, 9/10 M3 module genes cluster coherently in the lower-left quadrant confirming that disruption of the WT SpxA1:SpxA2 ratio by loss of either paralog collapses the cooperative BAC-induced expression program (Figure 5A, green dots). M4 module genes cluster in the upper-right under both conditions, consistent with their stoichiometry-independent constitutive SpxA2-dependent repression pattern (Figure 5A-B, purple dots). M1 module genes show a characteristic left-shift on the x-axis reflecting SpxA1-dependent maintenance, with *sodA* and *endoS* as the most extreme examples (Figure 5A-B, red dots). Under hNP-1, the M3 cooperative cluster shifts toward y-axis dominance indicating that defensin challenge engages a more SpxA2-dependent regulatory program at these loci while the overall module structure is preserved (Figure 5B, green dots). The coherent module-to-quadrant mapping across both stress conditions confirms that SpxA1:SpxA2 stoichiometry is a primary determinant of GAS transcriptional regulatory tone under cell envelope stress, with the nature of the stress signal modulating the magnitude and paralog-specificity of module-level responses.

**Figure 5.**
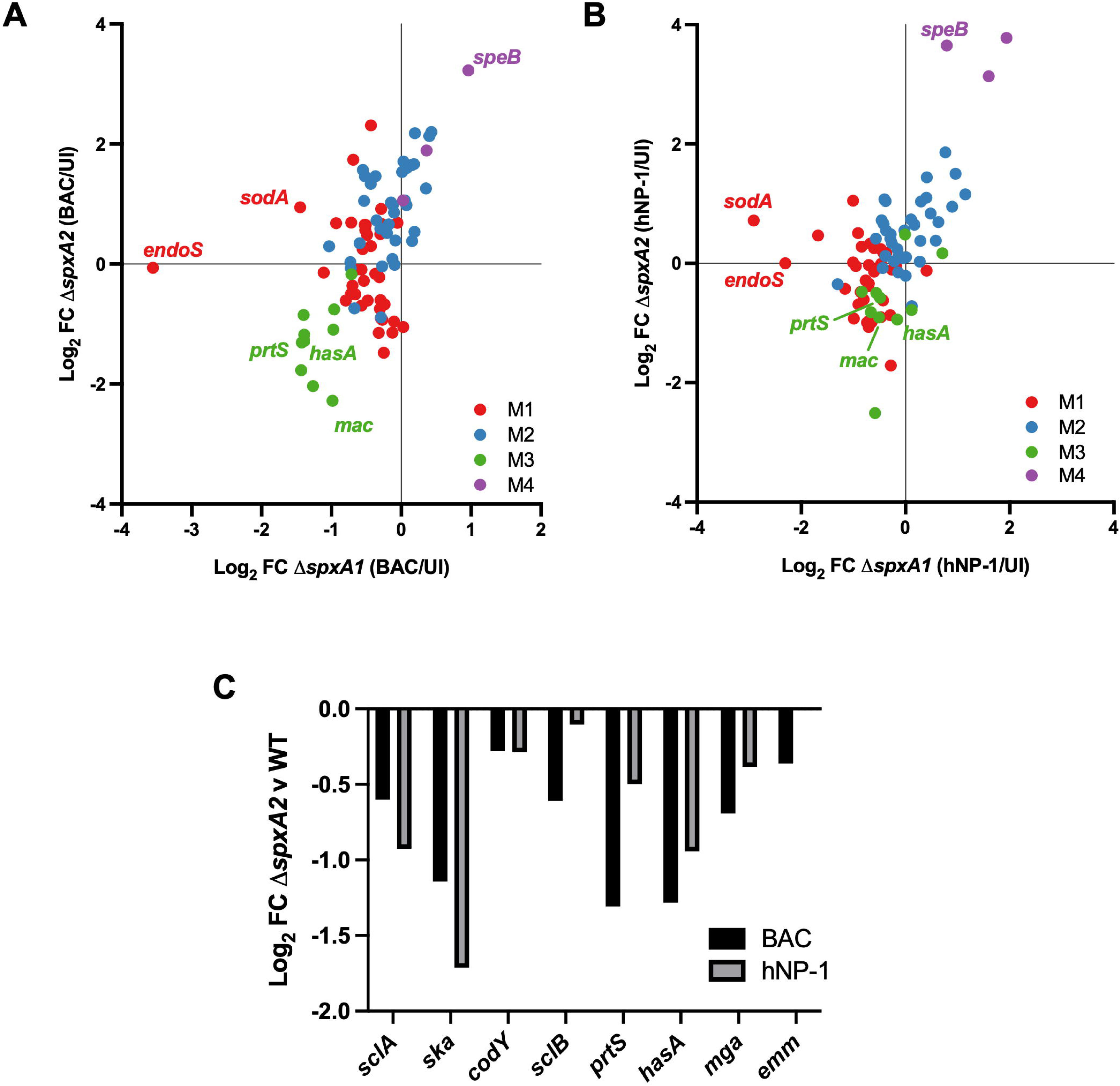
Condition-dependent paralog divergence revealed by direct Δ*spxA1*/ Δ*spxA2* transcriptional comparison. NanoString gene expression data from Figure 4 replotted as direct paralog comparisons. (**A**) Scatter plot comparing BAC-induced log_2_FC in Δ*spxA1* (x-axis) and Δ*spxA2* (y-axis). Coherent module-to-quadrant mapping confirms that SpxA1:SpxA2 stoichiometry drives module membership. (**B**) Scatter plot as in panel A comparing hNP-1-induced log_2_FC in Δ*spxA1* (x-axis) and Δ*spxA2* (y-axis). Overall module structure is preserved relative to BAC with reduced Δ*spxA1* displacement of M3 genes, indicating that defensin challenge engages a more SpxA2-dominant regulatory program at cooperative module loci. For A and B, each point represents one NanoString target gene, colored by module. (**C**) Bar chart showing NanoString log_2_FC of Δ*spxA2* vs. WT MGAS10870 for validated CovR-regulated target genes under BAC (black) and hNP-1 (gray) conditions. Genes are grouped by CovR binding motif following the framework of Horstmann et al. 2023 (18): Extended motif targets (*sclA*, *ska*, *codY*, *sclB*, *prtS*, *hasA*; left of dashed line) are subject to repression by CovR; ATTARA-only targets (*mga*, *emm*; right of dashed line) are subject to activation by CovR. Full NanoString DGE results are in Supplemental Table ST4.

### SpxA2 modulates transcription of CovR-regulated virulence determinants

We also assessed with greater scrutiny whether SpxA2 loss affected expression of validated CovR-regulated targets (18) using the NanoString dataset. CovR-regulated targets were distributed across the four gene modules in a pattern consistent with their regulatory motifs (Table 2; Supplemental Table ST5): Extended binding motif targets (subject to CovR repression) *hasA*, *prtS*, *scpA*, and *mac-1* were assigned to M3 (cooperative stress induction); *codY*, *sclA*, and *sclB*, to group to M1 (SpxA1-dependent maintenance); and *sagA* and *sagB* to M2 (SpxA2-dependent repression). Canonical CovR ATTARA-only binding motif targets (subject to CovR induction) *mga* and *emm*, assigned to M1. The distribution of CovR-regulated targets across multiple modules reflects SpxA2 influence on CovR-dependent transcription through both direct stress-induced mechanisms (M3) and effects on baseline regulatory tone (M1, M2).

Loss of SpxA2 was associated with reduced expression (i.e., repression) of the majority of gene targets with the extended CovR binding motif under both BAC and hNP-1 conditions (Figure 5C). Under BAC, all six extended motif targets assessed (*scl2/sclA*, *ska*, *codY*, *scl1*/*sclB*, *prtS*, *hasA*) were downregulated in Δ*spxA2* vs. WT, consistent with increased CovR-mediated repression at these promoters in the absence of SpxA2. Under hNP-1 a more pronounced repression was observed in Δ*spxA2* in *ska* (−1.71) and *sclA* (−0.93) compared to BAC (Figure 5C; Supplemental Table ST5). The extended motif target *sagA* was an exception [elevated in Δ*spxA2* under both BAC (log_2_FC = +1.26) and hNP-1 (log_2_FC = +0.95) (Supplemental Table ST5)] consistent with its M2 assignment where SpxA2-dependent repression is predicted to operate independently of CovR phosphorylation state potentially reflecting dual regulatory inputs at the *sagA* promoter. The genes defined by the ATTARA binding motif, e.g., *mga* and *emm*, showed directionally consistent downregulation in Δ*spxA2*, most prominently under BAC (*mga* log_2_FC = −0.69; *emm* log_2_FC = −0.36) (Figure 5C). The *nga/slo* operon showed variable expression across conditions, consistent with competing CovR-dependent and paralog-antagonistic regulatory inputs and precluding a consistent directionality interpretation.

### SpxA2 modulates CovR promoter DNA occupancy in a promoter-dependent manner

The NanoString and RNA-seq analyses described above define a LiaR -> SpxA2 -> CovRS regulatory axis that selectively influences a subset of GAS virulence gene targets in response to AMP-mediated cell envelope stress. To define the direct molecular basis for SpxA2-dependent modulation of CovR-regulated transcription, we performed chromatin immunoprecipitation with exonuclease treatment (ChIP-exo) using a validated polyclonal antibody directed against the N-terminal domain of CovR (36) (see Methods) in MGAS10870 (WT), the isogenic deletion of *spxA2* (Δ*spxA2*), and an isogenic mutant, LiaS^Q146A^, in which LiaS lacks the ability to dephosphorylate LiaR and therefore maintains constitutively elevated *spxA2* transcription (13). ChIP-exo libraries demonstrated high quality, with fraction of reads in peaks (FRiP) scores of 0.21–0.24 across all nine samples, and characteristic strand-specific 5’ read doublets flanking CovR binding sites confirming efficient λ-exonuclease digestion (see Supplemental Methods). Peak calling using MACS3 identified 362 high-confidence CovR binding sites in WT, of which 88% overlapped with previously published CovR ChIP-seq binding sites in the same strain (18), validating our approach. Differential binding analysis using DiffBind with DESeq2 normalization identified 439 differentially bound sites between WT and Δ*spxA2* (FDR < 0.05), compared to only 76 sites between WT and LiaS^Q146A^ (Supplemental Figures S7 and S8). The substantially greater effect of SpxA2 loss (Δ*spxA2*) compared to SpxA2 overexpression (LiaS^Q146A^) on CovR occupancy indicates that the relationship between SpxA2 levels and CovR binding is not simply linear. Thus, it is likely that additional factors including promoter architecture and CovR phosphorylation state constrain the response to elevated SpxA2 beyond the WT condition. Genome-wide composite analysis of ChIP-exo signal centered on WT peak summits confirmed a dose-dependent relationship, with Δ*spxA2* showing the highest CovR occupancy (suggesting gene repression), followed by LiaSQ^146A^, and WT showing the lowest occupancy (Figure 6A).

**Figure 6.**
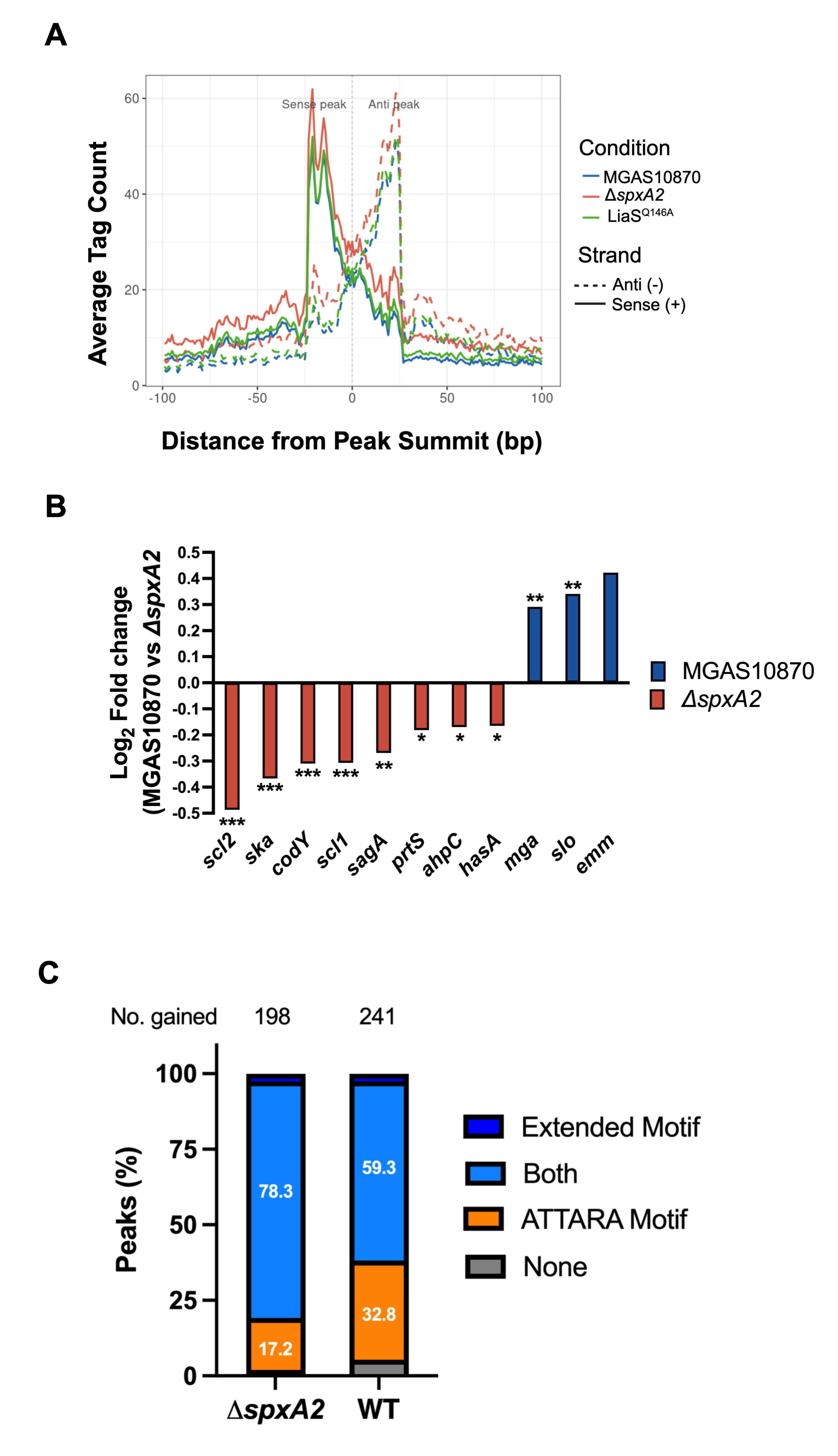
SpxA2 modulates CovR promoter DNA occupancy in a motif-dependent manner. CovR chromatin immunoprecipitation with exonuclease treatment (ChIP-exo) was performed in biological triplicate in MGAS10870, Δ*spxA2*, and LiaS^Q146A^ (SpxA2-OE) as described in Methods. (**A**) Composite ChIP-exo tag pileup centered on MGAS10870 peak summits (±100 bp) for MGAS10870 (blue), Δ*spxA2* (red), and LiaS^Q146A^ (green). Solid lines represent sense (+) strand signal; dashed lines represent anti (−) strand signal. Characteristic strand-specific 5′ read doublets flanking the peak summit confirm efficient λ-exonuclease digestion. Δ*spxA2* shows the highest average CovR occupancy, LiaS^Q146A^ intermediate, and MGAS10870 the lowest, consistent with a dose-dependent relationship between SpxA2 levels and CovR binding. (**B**) Log_2_FC in CovR binding occupancy (MGAS10870 vs. Δ*spxA2*) at known CovR-regulated virulence gene promoters. Negative values (red bars) indicate increased CovR occupancy in Δ*spxA2* (Δ*spxA2*-gained); positive values (blue bars) indicate increased CovR occupancy in MGAS10870 (MGAS10870-gained). Extended motif promoters susceptible to CovR-mediated repression (*scl2*, *ska*, *codY*, *scl1*, *sagA*, *prtS*, *ahpC*, *hasA*, *brnQ*) show increased CovR occupancy in Δ*spxA2*; ATTARA-only romoters susceptible to CovR-mediated activation (*mga*, *slo*, *emm*) show increased CovR occupancy in MGAS10870, consistent with SpxA2 facilitating CovR binding at these sites. Asterisks indicate statistical significance (* P < 0.05, ** P < 0.01, *** P < 0.001, FDR-corrected). (**C**) Motif composition at Δ*spxA2*-gained (n=198) and MGAS10870-gained (n=241) differentially bound peak sets. Peaks are classified by the presence of extended binding motif (WTWTTATAAWAAAAWNATDA) and/or ATTARA-only binding motif identified by FIMO motif analysis as described in Methods. Δ*spxA2*-gained peaks are significantly enriched for dimer-containing sequences (78.3% dimer only + 2.5% both motifs) relative to MGAS10870-gained peaks (59.3% + 2.5%; OR=1.76, P=0.0046, Fisher’s exact test). MGAS10870-gained peaks are significantly enriched for monomer-only sequences (32.8% vs. 17.2%; OR=2.35, P=0.00019, Fisher’s exact test), consistent with SpxA2 selectively antagonizing CovR dimer binding at extended motif promoters while facilitating CovR monomer binding at promoters with the ATTARA-only motif.

Among known CovR-regulated virulence factor-encoding genes, SpxA2 loss produced two distinct patterns of differential binding consistent with the extended and ATTARA binding motif framework established by Horstmann et al. 2023 (18). At eight promoters encoding major virulence factors, including *ska*, *hasA*, *scpC/prtS*, *sagA*, *scl1/sclB,* and *scl2/slcA*, CovR occupancy was significantly higher in Δ*spxA2* than WT (Figure 6B), consistent with SpxA2 normally antagonizing CovR binding at these extended motif loci. The increased CovR occupancy at promoters with the CovR extended binding motif in Δ*spxA2* is directly consistent with the reduced transcription of these targets observed in Δ*spxA2* under BAC and hNP-1 conditions (Figure 5A-B; Supplemental Table ST4 and ST5). That is, increased CovR-mediated repression at these promoters in the absence of SpxA2 provides the direct mechanistic basis for their transcriptional downregulation (i.e., repression). Conversely, at four promoters including those of *emm*, *mga*, and *nga/slo* (targets with the ATTARA-only binding motif) occupancy was significantly higher in WT than Δ*spxA2* (Figure 6B), suggesting SpxA2 promotes CovR binding at these sites. The reduced CovR occupancy at promoters with the canonical ATTARA motif in Δ*spxA2* similarly accounts for the downregulation of *mga* and *emm* transcription observed under BAC conditions (Figure 5A; Supplemental Table ST4 and ST5). These data establish that SpxA2 bidirectionally shapes the CovR-dependent transcriptional landscape de-repressing targets with the extended CovR binding motif while maintaining activation of targets with the ATTARA motif.

To investigate the mechanistic basis for these two motif classes of SpxA2-dependent CovR binding, we performed motif analysis using Find Individual Motif Occurrences (FIMO) (37) of the MEME suite to scan differentially bound peaks for the previously identified CovR extended (WTWTTATAAWAAAAWNATDA) and ATTARA-only binding motifs (18). Sites with increased CovR binding in Δ*spxA2* were significantly enriched for the extended binding motif compared to sites with decreased binding (66.7% vs. 53.1%, OR=1.76, p=0.0046, Fisher’s exact test). Furthermore, sites with decreased CovR binding in Δ*spxA2* were significantly enriched for sequences lacking the extended dimer motif (32.8% vs. 17.2%, OR=2.35, p=0.00019, Fisher’s exact test) (Figure 6C). These data indicate that SpxA2 selectively antagonizes CovR binding at promoters characterized by the extended dimer motif, while facilitating CovR binding at promoters characterized by the more restricted ATTARA motif. This motif-dependent bidirectional modulation of CovR-DNA occupancy by SpxA2 provides the direct molecular basis for the selective influence of SpxA2 on a subset of CovR-regulated virulence gene targets identified across the RNA-seq, proteomics, and NanoString analyses described above.

## Discussion

GAS encodes a complex regulatory network that coordinates virulence gene expression in response to host-derived signals, yet the mechanisms by which individual stress-sensing systems interact to reshape the broader virulence gene regulatory landscape have remained poorly defined. The present study establishes that SpxA2, acting downstream of the LiaFSR CES-sensing system, directly modulates CovR-DNA binding occupancy in a promoter motif-dependent manner to bidirectionally regulate CovR-controlled virulence gene expression. Integrating RNA-seq, DIA proteomics, NanoString-based transcriptional profiling across multiple stress conditions, and ChIP-exo, we further demonstrate that SpxA1 and SpxA2 are functionally non-redundant paralogs whose relative activity functions as a regulatory rheostat calibrating the overall virulence gene regulatory tone of the GAS cell (Figure 7). Our findings provide a mechanistically detailed example of a Gram-positive stress-responsive regulator directly shaping the DNA-binding activity of a master virulence regulator. The current study also establishes a framework for understanding how GAS and other Gram-positive pathogens integrate cell envelope stress signals into coordinated virulence gene regulatory changes to affect disease progression.

**Figure 7.**
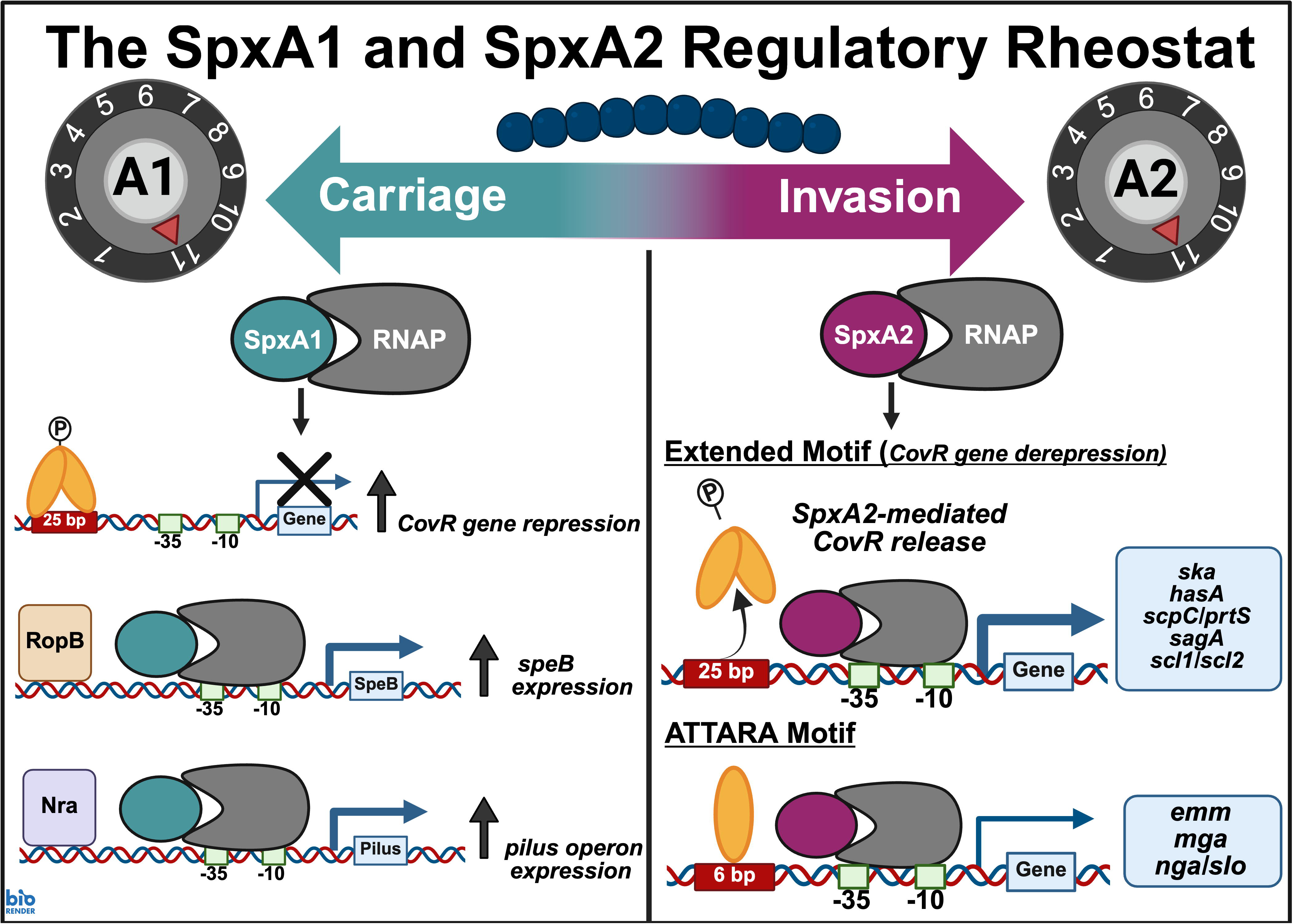
The GAS SpxA1 and SpxA2 Regulatory Rheostat. GAS exists along a continuum from asymptomatic colonization (carriage) to invasive disease marked by breaching of epithelial barriers and tissue dissemination. The current study supports a mechanistic framework in which these two states are governed by the relative SpxA1:SpxA2 stoichiometric balance, with each extreme characterized by a distinct transcriptional regulatory program. In the SpxA1-dominant carriage state, CovR-mediated repression of immune evasion genes is maintained, RopB drives SpeB expression, and pilus operon transcription is active – potentially through cooperative recruitment of RopB and Nra to target promoters via the SpxA1-RNAP complex, though the precise mechanism remains to be defined. Conversely, activation of LiaFSR by host-derived antimicrobial peptides directly increases SpxA2, shifting the stoichiometric balance toward SpxA2 dominance. In this state, SpxA2-RNAP engagement modulates CovR-DNA binding in a promoter motif-dependent manner – releasing phosphorylated CovR dimer repression at immune evasion gene promoters with the extended CovR binding motif while maintaining unphosphorylated CovR monomer activation at promoters with the ATTARA-only CovR binding motif – producing a virulence gene expression profile characteristic of invasive disease.

While the LiaFSR-dependent regulation of *spxA2* and the functional specialization of SpxA1 and SpxA2 as non-redundant regulatory paralogs are features conserved from *S. mutans* (14, 15, 21, 22), the present study defines their downstream consequences for CovRS-dependent virulence gene control as distinctly GAS contributions. The divergent regulon compositions of Δ*spxA1* (158 genes, oxidative stress-enriched) and Δ*spxA2* (419 genes, virulence and CovRS cross-regulation-enriched) confirm functional non-redundancy consistent with the division of regulatory labor described for *S. mutans* orthologs (21, 22). DIA proteomics provided the first demonstration that SpxA2 is a ClpXP substrate in GAS, extending prior observations from *B. subtilis*, *S. mutans,* and *S. aureus* (15, 19, 27), with differential ClpXP efficiency between paralogs consistent with their C-terminal sequence differences (15). The reciprocal accumulation pattern (i.e., SpxA2 elevated in Δ*spxA1* and SpxA1 trending elevated in Δ*spxA2*) suggests compensatory transcriptional upregulation when either paralog is absent, though the precise mechanism remains to be determined. C-terminal swap experiments further established that the SpxA2 functional domain is both necessary and sufficient for AMP tolerance independent of C-terminal sequence identity. Notably, GAS SpxA1 lacks the terminal acidic residues that confer ClpXP resistance in *S. mutans* SpxA1 (15), suggesting that functional domain rather than proteolytic turnover differences are the primary basis for paralog specialization in GAS. Together, these data support a model in which the SpxA1/SpxA2 balance is maintained by differential ClpXP proteolysis at baseline and shifted toward SpxA2 dominance upon LiaFSR activation.

High-resolution CovR mapping using ChIP-exo represents the first application of this technique in any streptococcal species, providing unprecedented genome-wide resolution of transcription factor-DNA binding occupancy in GAS. We applied this approach to characterize SpxA2 influence on CovR-DNA binding across strain backgrounds differing in SpxA2 levels. Importantly, the established framework may be extended to any GAS transcription factor beyond CovR revealing yet additional interconnected networks. SpxA2 loss produced 439 differentially bound CovR sites genome-wide compared to only 76 in the SpxA2-overexpressed condition, indicating that the effect of SpxA2 on CovR occupancy is not simply linear and that promoter architecture and CovR phosphorylation state constrain the response. At extended CovR motif promoters (18), SpxA2 loss increased CovR occupancy, consistent with SpxA2 normally antagonizing CovR dimer binding to de-repress immune evasion factors including *ska*, *hasA*, and *scl1/scl2*. Conversely, at ATTARA-only promoters potentially bound by CovR monomers, SpxA2 loss reduced CovR occupancy, consistent with SpxA2 facilitating monomer binding and maintaining activation of *emm* and *mga*. This bidirectional modulation of a single response regulator depending on phosphorylation state and promoter architecture is analogous to the *B. subtilis* Spx-ComA system (19, 20), but represents a more sophisticated regulatory logic not previously described for any Spx family member. While the motif composition data are fully consistent with the two structural CovR binding motifs defined by Horstmann et al. 2023 (18) in the same strain, formal TSS mapping will be required to fully resolve the spatial relationship between SpxA2-modulated binding sites and transcription initiation. Further, additional work is needed to define *emm* type-specific differences in SpxA2-CovR regulatory patterns and how such differences may contribute to strain-specific disease phenotypes.

NanoString-based transcriptional profiling of a targeted 116-gene panel revealed a structured four-module regulatory architecture that would not have been apparent from individual pairwise comparisons alone, validating the multi-condition clustering approach for dissecting paralog-specific regulatory logic. The four modules reveal that the SpxA1/SpxA2 stoichiometric ratio determines which transcriptional programs are engaged under a given stress condition. Most strikingly, M3 defines a cooperative program requiring simultaneous presence of both SpxA1 and SpxA2 for BAC-induced expression of CovR-regulated immune evasion targets – a program that collapses when either paralog is absent and exists only at the WT stoichiometric ratio under cell envelope stress. Together the four modules (Table 2) demonstrate that SpxA1 and SpxA2 influence multiple transcription factors beyond CovR, and that the regulatory tone of the GAS cell is a direct function of the SpxA1/SpxA2 stoichiometric balance set by LiaFSR-mediated induction, ClpXP-mediated proteolysis, and the nature of the prevailing host-derived stress signal.

The convergence of transcriptomic, proteomic, and NanoString data establishes SpxA1 as a primary determinant of the oxidative stress regulatory tone in GAS, operating in parallel to the SpxA2-dependent virulence gene regulatory program. Similar to *S. mutans* (22), redox homeostasis is maintained by SpxA1 in GAS revealed through the transcriptome (*sodA* and *nox.1* dysregulation) and proteomics (altered levels of DpR, GPr, PerR, and Rex). Reductions in DpR and GPr without corresponding transcript changes indicate indirect regulatory mechanisms, with SpxA1-mediated alteration of PerR protein levels (32) as the most likely candidate. The significant elevation of Rex in Δ*spxA1* provides additional evidence of a broad NAD^⁺^/NADH redox network shift consistent with loss of Nox.1 activity. Collectively, SpxA1 maintains an interconnected redox regulatory network whose disruption propagates to affect multiple redox-sensing transcription factors that mirrors the role of SpxA2 in maintaining CovR-dependent virulence gene regulatory tone. The functional consequences of SpxA1-dependent oxidative stress regulatory tone for GAS pathogenesis during host interaction are the subject of ongoing investigation.

The regulatory framework defined here provides a mechanistic basis for understanding the transition of GAS between asymptomatic carriage and invasive infection through the SpxA1:SpxA2 stoichiometric balance (Figure 7). In an SpxA1-dominant state, GAS maintains a transcriptional profile consistent with a carrier phenotype characterized by enhanced epithelial colonization (38, 39) (Figure 7). This model is directly supported by our prior work demonstrating that the naturally occurring carriage-associated LiaS^R135G^ variant produces a phenotype consistent with constitutively low SpxA2, leading to increased pilus production and enhanced epithelial adherence (12, 38). The current study supports the prior work by identifying the pilus operon (via *cbp* and *tee3*) as the likely targets of SpxA1/SpxA2 regulatory interaction given that *cbp* and *tee3* are M4 module targets that are elevated in expression in the absence of SpxA2. At the opposite end of the spectrum, LiaFSR activation by host-derived AMPs (i.e., hNP-1) shifts the balance toward SpxA2 dominance producing a transcriptional state consistent with immune evasion and invasion. Critically, that several attempts failed to produce a *spxA1*/*spxA2* double deletion is consistent with this framework. That is, complete loss of both paralogs would produce simultaneous collapse of oxidative stress defense, immune evasion gene maintenance, and CovR-dependent virulence gene regulatory capacity producing a regulatory paralysis incompatible with bacterial fitness. The SpxA1:SpxA2 rheostat thus emerges as a fundamental determinant of GAS lifestyle, with the LiaFSR system serving as the host-responsive molecular switch that shifts stoichiometric balance in response to the antimicrobial peptide environment of the human infection site. The methodological and conceptual framework established here provides the foundation for directly testing how SpxA1/SpxA2 regulatory balance shapes GAS-host interactions and has direct implications for understanding how this and other Gram-positive pathogens cause disease in humans.

## Materials and Methods

### Bacterial Strains and Culture Conditions

The GAS strain used in this study was MGAS10870, a serotype M3 strain originally isolated in ON, Canada, in 2002 from a patient with a soft tissue infection that represents the wild-type or parental strain (40). All strains used in this study are listed in Supplemental Table ST6. GAS strains were grown in Todd-Hewitt broth containing 0.2% (wt/vol) yeast extract (THY broth, DIFCO Laboratories), on THY agar, or Trypticase soy agar (TSA II) containing 5% sheep blood agar (Becton Dickinson). For all in vitro assays unless otherwise indicated, cultures were grown overnight without shaking at 37°C with 5% CO2 in THY broth to the required culture density. For RNA-seq and DIA proteomics, cultures were grown to exponential phase (OD_600_ 0.9–1.0). For NanoString studies, cultures were grown to early logarithmic phase (OD_600_ 0.4–0.5) prior to treatment with bacitracin (1 μg/mL; Sigma-Aldrich), hNP-1 (5 μg/mL; Bachem), or vehicle (uninduced; UI) for one additional hour before harvest.

### Generation of Isogenic Mutants

The primers used in this study are listed in Supplemental Table ST7. The isogenic mutants lacking either *spxA2* or *clpX* were previously published in Sanson et al. (12). The isogenic mutant with altered pattern of LiaR phosphorylation, LiaS^Q146A^ [generated in MGAS10870 background by the in-frame replacement of nucleotides in the *liaS* (SpyM3_1367) coding sequence resulting in glutamine to alanine at position 146 (Q146A)] was previously published in Vega et al. (13). The isogenic mutant lacking *spxA1* was made by in-frame insertional inactivation with a kanamycin resistance gene (*aph*) as previously described (41, 42). For both the promoter and C-terminal “swapped” *in-trans* mutant strains, constructs were synthesized (Genewiz; Azenta Life Sciences) and inserted into pLZ12 (43) using *Bam*HI and *Xho*I restriction. Electrocompetent cells of Δ*spxA2* were transformed with these plasmids as previously described (44).

### RNA-sequencing Analysis

RNA-seq was performed as previously described (45). Briefly, GAS cultures were grown in THY broth in biological triplicate to exponential phase, harvested into RNAprotect (Qiagen), and RNA isolated using the RNeasy mini kit (Qiagen). cDNA libraries were prepared using the Script-Seq Complete Kit (Bacteria; Epicentre/Illumina) and sequenced on a NextSeq 2000 (average ∼62 million 150 bp paired-end reads per replicate). Reads were mapped to the MGAS10870 reference genome (GenBank CP067090) using CLC Genomics Workbench v23 (Qiagen). Differential gene expression analysis was performed using DESeq2 (46) with Benjamini-Hochberg correction; thresholds of *P-*adj < 0.05 and |log_2_FC| ≥ log_2_(1.5) were applied. Prophage regions and deletion targets were excluded from analysis. Full quality control metrics, mapping parameters, and differential expression results are provided in Supplemental Methods and Supplemental Table ST1.

### NanoString nCounter Analysis

NanoString-based gene expression profiling was performed using a custom 116-gene codeset designed against the MGAS10870 *emm3* reference genome (GenBank CP067090). RNA (100 ng per sample) from cultures grown under UI, BAC, or hNP-1 challenge conditions (see Bacterial Strains and Culture Conditions) was hybridized according to the nCounter Gene Expression Assay manufacturer’s protocol (Bruker Spatial Biology) and processed at the Vanderbilt Technologies for Advanced Genomics (VANTAGE) core facility. Differential expression analysis was performed using the limma empirical Bayes framework (34, 35) on nSolver-normalized counts. One biological replicate (Δ*spxA1* hNP-1, replicate A) was excluded as a PCA outlier. Full details of codeset design, normalization and quality control, cross-platform validation with RNA-seq, and gene module clustering analysis are provided in Supplemental Methods. Complete differential expression results are provided in Supplemental Table ST4.

### Antimicrobial and Antimicrobial Peptide tolerance

Tolerance to bacitracin (BAC) and human neutrophil peptide-1 (hNP-1) was assessed using virtual colony counts (CFU_V_) as previously described (13). Briefly, GAS strains were grown to OD_600_ 0.45–0.55 (∼10^8^ CFU/mL), diluted 20-fold in sodium phosphate buffer (10 mM, pH 7.4), and exposed to 2x final concentrations of AMP or vehicle in a 96-well format for 90 minutes at 37°C with 5% CO_2_. Following treatment, cultures were diluted into fresh THY and outgrowth was monitored by OD_600_ over 20 hours on a Synergy H1 plate reader (BioTek). CFU_V_ values were interpolated from calibration curves as previously described (13). Full methodological details including calibration curve generation and CFU_V_ calculation are provided in Supplemental Methods.

### DIA Proteomics and Analysis

GAS cultures were grown in THY broth in biological triplicate to exponential phase (OD_600_ 0.9–1.0). Cell lysates were prepared in PBS with cOmplete EDTA-free Protease Inhibitor Cocktail (Roche) and 1.4 mM diamide (Sigma-Aldrich) by mechanical lysis using the FastPrep-24 (MP Biomedicals). Protein concentration was determined by Bradford assay (Bio-Rad). Sample processing, trypsin digestion, and DIA data acquisition were performed by the Mass Spectrometry Research Center, Vanderbilt University Medical Center. Peptides were resuspended in 0.2% FA to a concentration of 200 ng/μL. Peptide concentrations were estimated using a Nanodrop 2000 Spectrophotometer (Thermo Scientific), and samples were adjusted to 50 ng/μL. 0.015% n-dodecyl-β-D-maltoside was added to diaPASEF samples. DiaPASEF data were acquired on 150 ng of peptides using a 30-min aqueous-to-organic gradient method delivered via a nanoELUTE2 360 μm outer diameter × 100 μm internal diameter column packed with 20 cm of 3 μm C18 reverse phase material (Jupiter, Phenomenex) coupled to a timsTOF HT or Ultra2 instrument (Bruker) diaPASEF data were collected in 12 PASEF ramps from 0.75 to 1.3 1/*k* _0_ covering 350 to 1250 *m*/*z* via variable windows ranging in size from 12.27 to 122.81 Th with 50 ms accumulation time. Raw data were processed using Spectronaut V20 (Biognosys) *Streptococcus pyogenes* MGAS315 proteome (UniProt proteome UP000000564). Differential abundance analysis was performed in R (v4.5.3) using the limma package with empirical Bayes variance moderation and Benjamini-Hochberg correction; full details are provided in Supplemental Methods. Complete proteomics results are provided in Supplemental Table ST2.

### ChIP-exo

ChIP-exo was performed by The Cornell Institute of Biotechnology BRC Epigenomics Core Facility using a validated polyclonal antibody directed against the N-terminal domain of CovR, as previously described (36). Three biological replicates of MGAS10870, Δ*spxA2*, and LiaS^Q146A^ were grown in THY to mid-exponential phase (OD_600_ ∼0.45), cross-linked with 1% formaldehyde, and processed for ChIP-exo. Libraries were sequenced on an Illumina platform (75-nucleotide single-read format) and reads were mapped to the MGAS10870 reference genome (GenBank NZ_CP067090.1) using CLC Genomics Workbench v21. FRiP scores of 0.21–0.24 confirmed high ChIP enrichment quality across all nine samples (Supplemental Figure S10). CovR binding peaks were called using MACS3 and differential binding was assessed using DiffBind v3.20 with DESeq2 normalization; sites with FDR < 0.05 were considered significantly differentially bound. Motif analysis was performed using FIMO v5.5.9 from the MEME Suite (37). Full bioinformatic pipeline details including peak calling parameters, differential binding analysis, motif analysis, and gene annotation are provided in Supplemental Methods.

### Statistical analyses

Statistical analysis was performed using GraphPad Prism version 10, or R (v4.5.3). For parametric data with two groups, a one-tailed Student’s t test was used. All data analyzed in this work were derived from at least three fresh biological replicates.

## Data and Code Availability

All R and Python analysis code is available at https://github.com/ar-flores/SpxA1-SpxA2-MultiOmic-GAS. Raw sequencing data (RNA-seq, ChIP-exo) and NanoString normalized count data have been deposited in the NCBI Gene Expression Omnibus (GEO) under accession numbers GSE334312 (RNA-seq), GSE334313 (ChIP-exo), and GSE334410 (NanoString). Raw proteomics data have been deposited to the ProteomeXchange Consortium via the PRIDE partner repository with the dataset identifier PXD079489.

## Acknowledgments

This work was supported in part by NIAID R01AI125216 to A.R.F. and NIAID R21AI177931 to S.A.S. Vanderbilt Technologies for Advanced Genomics (VANTAGE) core facility is supported in part by the Vanderbilt Ingram Cancer Center (P30 CA68485), the Vanderbilt Vision Center (P30 EY08126), the NIH/NCRR (G20 RR030956) and an NIH High-End Equipment Grant (S10OD025092).

## Author Contributions

G.A.M., L.A.V., and M.S.I. performed experiments, analyzed data, and drafted the manuscript. K.H. performed experiments. W.H.M. generated mass spectrometry data. N.H. and S.A.S. performed ChIP-exo data analysis and edited the manuscript. J.A.G. contributed to conceptualization, performed experiments, and edited the manuscript. A.R.F. conceived and supervised the study, acquired funding, and edited the manuscript. All authors reviewed and approved the final manuscript..

