## Supplemental Methods for "SpxA1 and SpxA2 Function as a Stoichiometry-Dependent Regulatory Rheostat Governing Virulence Gene Expression in Group A Streptococcus"

**RNA-sequencing — detailed methods**

*Library preparation and sequencing*

GAS cultures were grown in THY broth (biological triplicate) to exponential phase (OD_600_ 0.9-1.0). Cells were harvested by centrifugation following addition of RNAprotect bacteria reagent (Qiagen). RNA was isolated and purified using the RNeasy mini kit (Qiagen) according to the Gram-positive bacteria protocol. Quantity and quality were determined using a Nanodrop (ThermoFisher) and Agilent 4200 TapeStation system. RNA purification, rRNA depletion, and adapter-tagged cDNA library preparation were performed using the Script-Seq Complete Kit (Bacteria; Epicentre/Illumina). cDNA libraries were sequenced on a NextSeq 2000 (Illumina); average library size was ~62 million (range 48–77 million) 150 bp paired-end reads per replicate. Following onboard adapter trimming, reads were mapped to the MGAS10870 reference genome (GenBank CP067090) using CLC Genomics Workbench v23 (Qiagen). Gene-level read counts, TPM, and RPKM were exported for all 1,888 annotated CDS features.

*Quality control*

Library sizes ranged from 16.4 to 20.0 million reads per sample. Principal component analysis of variance-stabilized transformed counts showed clean separation of the three strain groups on PC1 (44.3%) and PC2 (27.6%), explaining 72.9% of total variance. All biological replicates clustered within their respective strain groups with no outliers identified. DESeq2 dispersion estimates showed the expected decreasing mean-dispersion relationship, confirming appropriate model fit for the negative binomial distribution. QC figures including principal component analysis (PCA), MA plots, and sample-to-sample Euclidean distances, are provided in Supplemental Figure S1.

*Differential gene expression analysis*

Differential expression analysis was performed using DESeq2 (R/Bioconductor, version 1.48.2) (1) on raw count data from all nine samples. Genes with fewer than 10 total counts across all samples were excluded prior to analysis (1,799 of 1,888 genes retained). Size factors were estimated using the median ratio method (range 0.953–1.248). Variance-stabilized transformation (vst, blind = TRUE) was used for quality control visualization. Differential expression was assessed for Δ*spxA1* vs. WT and Δ*spxA2* vs. WT using the Wald test with Benjamini-Hochberg correction. Significance thresholds of *P*-adj < 0.05 and |log_2_FC| > log_2_(1.5) were applied. Prophage regions (C9Q_00070–00240; C9Q_00520–00595; C9Q_01370–01405; C9Q_02650–02935; C9Q_05500–05755; C9Q_06060–06365; C9Q_06545–06800; C9Q_07120–07360; 332 total locus tags) and deletion targets (*spxA1*, C9Q_04800; *spxA2*, C9Q_09105) were excluded. Full DESeq2 results for both contrasts are provided in Supplemental Table ST1.

**NanoString nCounter analysis — detailed methods**

*Codeset design*

A custom NanoString nCounter CodeSet was designed to simultaneously quantify 119 transcripts comprising 116 target genes and 3 endogenous control probes (*tuf*, C9Q_02490; *gyrA*, C9Q_04425; *guaB*, C9Q_09385). The codeset was developed against the *S. pyogenes* MGAS10870 (*emm3*; GenBank CP067090) and MGAS2221 (*emm1*; GenBank CP043530) reference genomes. Target genes were selected to represent the GAS transcriptome including virulence factor-encoding genes, transcriptional regulators, stress response factors, and metabolic enzymes. Five probes targeting *emm1*-specific genes absent from the *emm3* genome (*rofA*, *cpa*, *prtF*, *srtB*, *sic1*) were retained in the codeset; cross-hybridization signal from these probes in the *emm3* background is expected and noted in figure legends. The complete probe list is provided in Supplemental Table ST3.

*Normalization and quality control*

Raw NanoString count data were processed and normalized using nSolver Analysis Software (version 4.0, Bruker Spatial Biology). Quality control metrics assessed for each sample included: field of view counted (≥75% required), binding density (0.05–2.25), positive control linearity (R² ≥ 0.95), and limit of detection. No samples failed technical quality control. Normalization was performed sequentially: (1) positive control normalization for lane-to-lane hybridization efficiency; (2) background subtraction using the mean plus two standard deviations of eight negative control counts; and (3) endogenous normalization using the geometric mean of the three control probes. Normalized count data are deposited in GEO under accession number GSE334410.

*Differential gene expression analysis*

Differential expression analysis was performed using the limma empirical Bayes framework (R/Bioconductor limma, version 3.64.3) (2, 3) applied to log_2_-transformed nSolver-normalized counts. A linear model was fit using a cell-means parameterization with a single group factor encoding all strain × condition combinations (9 levels: WT, Δ*spxA1*, and Δ*spxA2* each under UI, BAC, and hNP-1). Twelve pairwise contrasts were defined: six within-strain comparisons (BAC vs. UI and hNP-1 vs. UI within each strain) and six cross-strain comparisons (each deletion mutant vs. WT under matched conditions). Empirical Bayes moderation was applied using the eBayes function. Differential expression thresholds of |log_2_FC| > log_2_(1.5) and unadjusted *P* < 0.05 were applied as working criteria; Benjamini-Hochberg FDR-adjusted *P*-values are provided in Supplemental Table ST4. One biological replicate (Δ*spxA1* hNP-1, replicate A) was identified as a global expression outlier by PCA (Supplemental Figure S4) and hierarchical sample clustering (Supplemental Figure S5) in the absence of technical quality control flags and was excluded from all analyses.

*Cross-platform validation with RNA-seq*

Cross-platform expression concordance was assessed at three levels. First, NanoString log_2_-normalized counts in WT uninduced conditions were correlated with RNA-seq log_2_(TPM + 1) values across 86 mapped panel genes (Pearson r = 0.781, R^2^ = 0.611; Supplemental Figure S9A). Second, NanoString log_2_FC values from limma analysis were compared to DESeq2 log_2_FC values for Δ*spxA1* vs. WT (r = 0.666, n = 90) and Δ*spxA2* vs. WT (r = 0.687, n = 90; Supplemental Figure S9B–C). Third, among genes identified as differentially expressed by NanoString analysis, directional concordance with DESeq2 log_2_FC was 82.9% for Δ*spxA1* vs. WT and 85.7% for Δ*spxA2* vs. WT. The RNA-seq and NanoString experiments were performed independently with different biological replicates; concordance reflects cross-platform biological reproducibility rather than technical replication.

*Gene module analysis*

To identify gene sets with shared regulatory logic, log_2_FC profiles were compiled across all 12 valid contrasts. Probes for *spxA1* and *spxA2* were excluded as their differential expression is expected by design. Genes with |log_2_FC| > log_2_(1.5) in at least two contrasts were retained for clustering (86 of 114 probes), and their log_2_FC values were assembled into an 86 x 12 matrix. Unsupervised hierarchical clustering was performed using Euclidean distance with complete linkage (hclust, R). Silhouette analysis identified k = 2 as the statistical optimum; a biologically motivated k = 4 solution was selected based on functional coherence of the resulting modules, mapping of ChIP-exo-validated CovR target genes to specific modules, and stability across random seeds (set.seed(42)). The k = 5 solution was evaluated and rejected as it produced only a magnitude gradient split within one module. Module assignments, mean log_2_FC profiles, and the full log_2_FC heatmap are provided in Supplemental Table ST5 and Supplemental Figure S5.

**Antimicrobial and antimicrobial peptide tolerance assays – detailed methods**

Seed cultures were grown from single colonies in 10 mL THY overnight. Cultures (500 μL) were diluted into 10 mL THY and grown to OD_600_ 0.45–0.55 (~10^8^ CFU/mL). A 20-fold dilution in sodium phosphate buffer (10 mM, pH 7.4) yielded ~5 × 10^6^ CFU/mL. AMPs were diluted to 2x final concentration in buffer and aliquoted (50 μL) in triplicate in a 96-well plate. GAS cell suspension (50 μL) was added to each well (~5 × 10^5^ CFU/well) and incubated for 90 minutes (37°C, 5% CO2). Following treatment, 100 μL 2x THY was added to each well. A 20 μL aliquot from each well was transferred to 180 μL pre-warmed THY for outgrowth monitoring by OD_600_ over 20 hours on a Synergy H1 microplate reader (BioTek; 37°C, 5% CO2). A 10-fold dilution series of untreated wells was plated on TSA-II/SB for CFU quantification to determine starting concentration C′0. OD_600_ measurements were normalized by subtracting blank readings. Calibration curves were generated by plotting threshold crossing times (ΔOD600 = 0.02) against log(C′0); linear regression r^2^ values ranged from 0.98 to 0.999. CFU_V_ values for treated samples were interpolated from strain-specific calibration curves. Outlier CFU_V_ values were identified and removed using Prism (GraphPad; Q = 5%). Survival ratio was calculated as CFU_V_ (AMP-treated) / C′0.

**DIA proteomics – differential abundance analysis**

All downstream analyses were performed in R (v4.5.3). LFQ intensities were log_2_-transformed and proteins were retained for analysis if ≥ 2 of 3 biological replicates had valid (non-zero) intensity values in at least one strain group, yielding 1,269 proteins for differential abundance testing. Pairwise differential abundance between each isogenic mutant strain and the parental MGAS10870 was assessed using the limma package (v 3.64.3) with empirical Bayes variance moderation. Contrast matrices were defined for Δ*spxA1* vs. WT and Δ*spxA2* vs. WT. *P*-values were adjusted for multiple testing using the Benjamini-Hochberg method. Proteins were considered significantly changed at |log_2_FC| ≥ log_2_(1.5) and adjusted *P*-value < 0.05, thresholds consistent with those applied in RNA-seq and NanoString analyses.

**ChIP-exo – detailed bioinformatic pipeline**

*Read processing and peak calling*

Raw sequencing reads were quality filtered, trimmed, and aligned to the MGAS10870 reference genome (GenBank NZ_CP067090.1) using CLC Genomics Workbench v26 (Qiagen). Replicate BAM files were merged per condition and indexed using SAMtools v1.13 (4). All samples achieved 100% mapping rates with 98.9–99.2% properly paired reads. Duplicate reads were removed by the sequencing core prior to data delivery. CovR binding peaks were called from merged BAM files using MACS3 (v3.0.4) with parameters: genome size 1,840,000 bp, --nomodel, --extsize 50, --shift -25, --keep-dup all, P-value threshold 1x10^-10^. Peaks with score > 5,000 were retained as high-confidence binding sites (362, 425, and 355 peaks for MGAS10870, Δ*spxA2*, and LiaS^Q146A^, respectively). A more permissive threshold of score > 3,000 was applied for DiffBind analysis.

*Differential binding analysis*

Differential CovR binding was assessed using DiffBind v3.20 (5) in R v4.5.3. Individual replicate BAM files were counted at 700 consensus peak regions using summit-centered 150 bp windows (summits = 75). Library-size normalization was applied using DESeq2 (1). Differential binding was assessed using DESeq2 with WT as the reference condition; sites with FDR < 0.05 were considered significantly differentially bound. All pairwise comparisons were performed: MGAS10870 vs. Δ*spxA2* (439 sites), MGAS10870 vs. LiaSQ146A (76 sites), and LiaSQ146A vs. Δ*spxA2* (378 sites).

*Strand-specific tag pileup and composite plots*

Strand-specific tag pileup was performed using ScriptManager v0.14 (Pugh lab, Cornell University; https://github.com/CEGRcode/scriptmanager). BAM files were converted to scIDX format using the bam-to-scidx function requiring proper mate pairs and outputting read 1 only to capture 5′ exonuclease stop positions. Tag pileup was generated centered on MACS3 peak summits ±100 bp separately for sense and anti-strands. Composite plots represent the average tag count across all 362 WT peak summits per position per strand for each condition.

*Motif analysis*

CovR binding motif occurrence was assessed using FIMO v5.5.9 from the MEME Suite (6). Two CovR binding motifs identified in MGAS10870 by Horstmann *et al.* (2023) (7) were used as queries: the dimer motif (WTWTTATAAWAAAAWNATDA) and the monomer motif (ATTARA). FIMO was run with a *P*-value threshold of 0.001 against sequences extracted from KO-gained (n = 198) and WT-gained (n = 241) differentially bound peak classes. Genomic sequences were extracted using bedtools getfasta v2.30.0 (8) from the reference genome FASTA generated from the NCBI GenBank file using Biopython v1.87. The proportion of sequences containing each motif class was compared between peak groups using Fisher's exact test in R v4.5.3. De novo motif discovery was performed using MEME v5.5.9 with parameters: -dna, -mod zoops, -revcomp, -nmotifs 3, -minw 10, -maxw 25 for KO-gained peaks and -minw 4, -maxw 15 for WT-gained peaks.

*Gene annotation*

Gene annotations were derived from the NCBI RefSeq reannotated MGAS10870 genome (NZ_CP067090.1). A BED file of gene coordinates was generated from the GenBank file using Biopython v1.87, capturing both gene and pseudogene feature types and prioritizing gene names over RefSeq locus tags (C9Q_RS prefix) where available. Known CovR binding sites were cross-referenced to updated locus tag nomenclature as several genes from Horstmann *et al.* (2023) (7) have been reannotated in the current MGAS10870 genome.

**Data Accessibility**

All R and Python analysis code is available at https://github.com/ar-flores/SpxA1-SpxA2-MultiOmic-GAS. Raw sequencing data (RNA-seq, ChIP-exo) and NanoString normalized count data have been deposited in the NCBI Gene Expression Omnibus (GEO) under accession numbers GSE334312 (RNA-seq), GSE334313 (ChIP-exo), and GSE334410 (NanoString). Raw proteomics data have been deposited to the ProteomeXchange Consortium via the PRIDE partner repository with the dataset identifier PXD079489.

**LIST OF SUPPLEMENTAL TABLES AND FIGURES**

| **TABLE #** | **FILE NAME** |
| --- | --- |
| **ST1** | Spx_mBio-ST1_RNAseq-DGE_05-24-2026 |
| **ST2** | Spx_mBio-ST2_DIA-LFQ_05-24-2026 |
| **ST3** | Spx_mBio-ST3_NS-codeset_05-24-2026 |
| **ST4** | Spx_mBio-ST4_NS-DGE_05-24-2026 |
| **ST5** | Spx_mBio-ST5_NS-Modules_05-24-2026 |
| **ST6** | Spx_mBio-supp-methods_05-25-2026 |
| **ST7** | Spx_mBio-supp-methods_05-25-2026 |

| **FIGURE #** | **FIGURE TITLE** |
| --- | --- |
| **S1** | RNA-seq quality control metrics |
| **S2** | C-terminal regions of SpxA1 and SpxA2 of GAS show similar amino acid (AA) composition to their *S. mutans* orthologs |
| **S3** | Global DIA proteomics differential abundance analysis |
| **S4** | NanoString nCounter principal component analysis |
| **S5** | NanoString nCounter sample-to-sample Pearson correlation heatmap |
| **S6** | NanoString gene module log_2_FC heatmap |
| **S7** | ChIP-exo differential CovR binding MA plot – WT vs. ∆*spxA2* |
| **S8** | ChIP-exo differential CovR binding MA plot – WT vs. LiaS^Q146A^ |
| **S9** | Cross-platform validation of NanoString nCounter against RNA-seq |
| **S10** | ChIP-exo library enrichment quality – Fraction of Reads in Peaks (FRiP) scores |

****all supplemental figures located in Spx_mBio-supp-methods_05-25-2026**

| **Table ST6.** Bacterial strains used in studies | | |
| --- | --- | --- |
| **Strain Name** | **Description** | **Reference** |
| MGAS10870 | Wild type serotype *emm3* strain background |  |
| *∆spxA2* | In-frame insertional inactivation of *spxA2* (*SpyM3_1799)* with a spectinomycin resistance cassette (*aad9*) | Sanson et al. (12) |
| *∆spxA1* | In-frame insertional inactivation of *spxA1* (*SpyM3_0885)* with a kanamycin resistance cassette (*aph*) | This study |
| *∆clpX* | in-frame insertional inactivation of *clpX* (*SpyM3_0604*) with a spectinomycin resistance gene (*aad9*) | Sanson et al. (12) |
| *∆spxA2::*pA2_A2 | *In trans* expression of SpxA2 from the *spxA2* promoter in the *∆spxA2* mutant background | This study |
| *∆spxA2::*pA1_A2 | *In trans* expression of SpxA1 from the *spxA2* promoter in the *∆spxA2* mutant background | This study |
| *∆spxA2::*pA2_A1 | *In trans* expression of SpxA2 from the *spxA1* promoter in the *∆spxA2* mutant background | This study |
| *∆spxA2::*A2_A1C | *In trans* expression of SpxA2 with the SpxA1 C-terminal region (115-134 AAs) from the *spxA2* promoter in the *∆spxA2* mutant background | This study |
| *∆spxA2::*A1_A2C | *In trans* expression of SpxA1 with the SpxA2 C-terminal region (115-134 AAs) from the *spxA2* promoter in the *∆spxA2* mutant background | This study |
| LiaS^Q146A^ | Substitution of Alanine in place of Glutamine at protein residue position 146 in LiaS in the *MGAS10870* background | Vega et al. (13) |

| **Table ST7.** Primers used in studies | | |
| --- | --- | --- |
| **Gene target** | **Sequence** | **Use** |
|  | AAACTCGAG**TTA**AAAAGAGCC | 3’ For to construct plZ12::pA2_A1 and plZ12::pA2_A1C to generate pA2_A1 and A2_A1C |
|  | CTTGGAAACTAAGGATCCAAA | 5’ reverse to construct plZ12::pA1_A2 to generate pA2_A1 |
| SpyM3_1799  (*spxA2)* | AAAAGGATCC**TTA**GAGTGCAGCACGTAAT | 3’ rev to construct plZ12::SpxA1_A2C to generate A1_A2C |
|  | GAAAGCCAAAACTTGGCTA | 3’ For to construct plZ12::SpxA2_A1C to generate A2_A1C |
|  | TAGCCAAGTTTTGGCTTTC | 5’ Rev to construct plZ12::SpxA2_A1C to generate *∆spxA2::*A2_A1C |
| SpyM3_0885  (*spxA1)* | GCAGCCAATTAAGCTACGCTC | 3’ flanking primer for *spxA1*::*aph* (3’ Fwd) |
|  | ATATTACTTATCCAAAGAAGCG | 5’ flanking primer for *spxA1*::*aph* (5’ Rev) |
|  | GTGGATAATAATAATTACCTGACATATCAGAAGAACTCGTCAAGAAGGCGATAG | 3’ spxa1 overlap primer for amp of kanamycin resistance gene, aph (3’ Fwd) |
|  | CTATCGCCTTCTTGACGAGTTCTTCTGATATGTCAGGTAATTATTATTATCCAC | 3’ spxa1-aph overlap primer for amplification of 3’ flanking region (3’ Rev) |
|  | GCGTGCAATCCATCTTGTTCAATCATATTTTTCCTCTATCAAATTTACTTACC | 5’ spxa1-aph overlap primer for amp of 5’ flanking region (5’ fwd) |
|  | GGTAAGTAAATTTGATAGAGGAAAAATATGATTGAACAAGATGGATTGCACGC | 5’ spxA1-aph overlap primer (5’ rev) |
|  | CGTTATAATGATGGGATCCAAA | 5’ reverse to construct plZ12::pA2_A1 and generate pA2_A1 |
|  | AAACTCGAGTTGCTAATAAAATC | 3’ For to construct plZ12::pA1_A2 and plZ12::pA1_A2C to generate pA1_A2 and A1_A2C |
|  | CAACTTCGTGTTTTACTAAC | 5’ Rev to construct Plz12::SpxA1_A2C to generate A1_A2C |
|  | GTTAGTAAAACACGAAGTTG | 3’ For to construct Plz12::SpxA1_A2C to generate A1_A2C |
|  | AAAAAAGGATCCTAAAAAAAGCTCTACAAGAGC | 3’ Rev to construct 3’ For to construct Plz12::SpxA2_A1C to generate A2_A1C |

**SUPPLEMENTAL FIGURES**

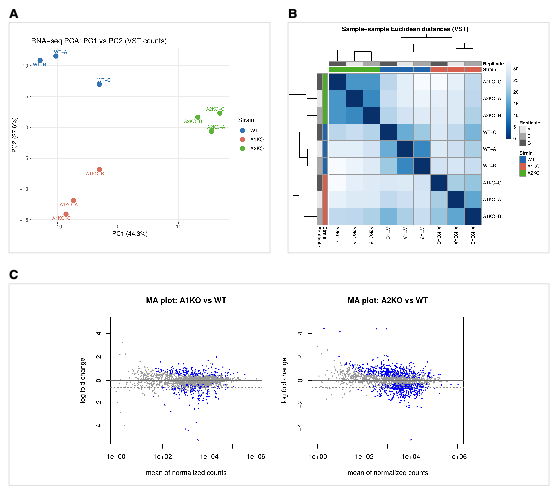

**FIGURE S1.** RNA-seq quality control metrics. (**A**) Principal component analysis (PCA) of variance-stabilized transformed (VST) RNA-seq counts across nine samples (WT, Δ*spxA1*, and Δ*spxA2*, biological triplicate). PC1 (44.3%) and PC2 (27.6%) account for 72.0% of total variance; replicates cluster tightly within strain groups confirming high inter-replicate reproducibility and clean separation of the three strain backgrounds. (**B**) Sample-to-sample Euclidean distance heatmap of VST counts. Color intensity reflects pairwise dissimilarity; within-strain distances (diagonal blocks) are substantially lower than between-strain distances, confirming distinct transcriptional profiles for each strain background. (**C**) MA plots for Δ*spxA1* vs. WT (left) and Δ*spxA2* vs. WT (right) showing log_2_ fold-change against mean normalized counts for all expressed genes. Blue dots indicate genes meeting significance thresholds (*P*-adj < 0.05, |log_2_FC| ≥ log_2_(1.5)); gray dots are non-significant.

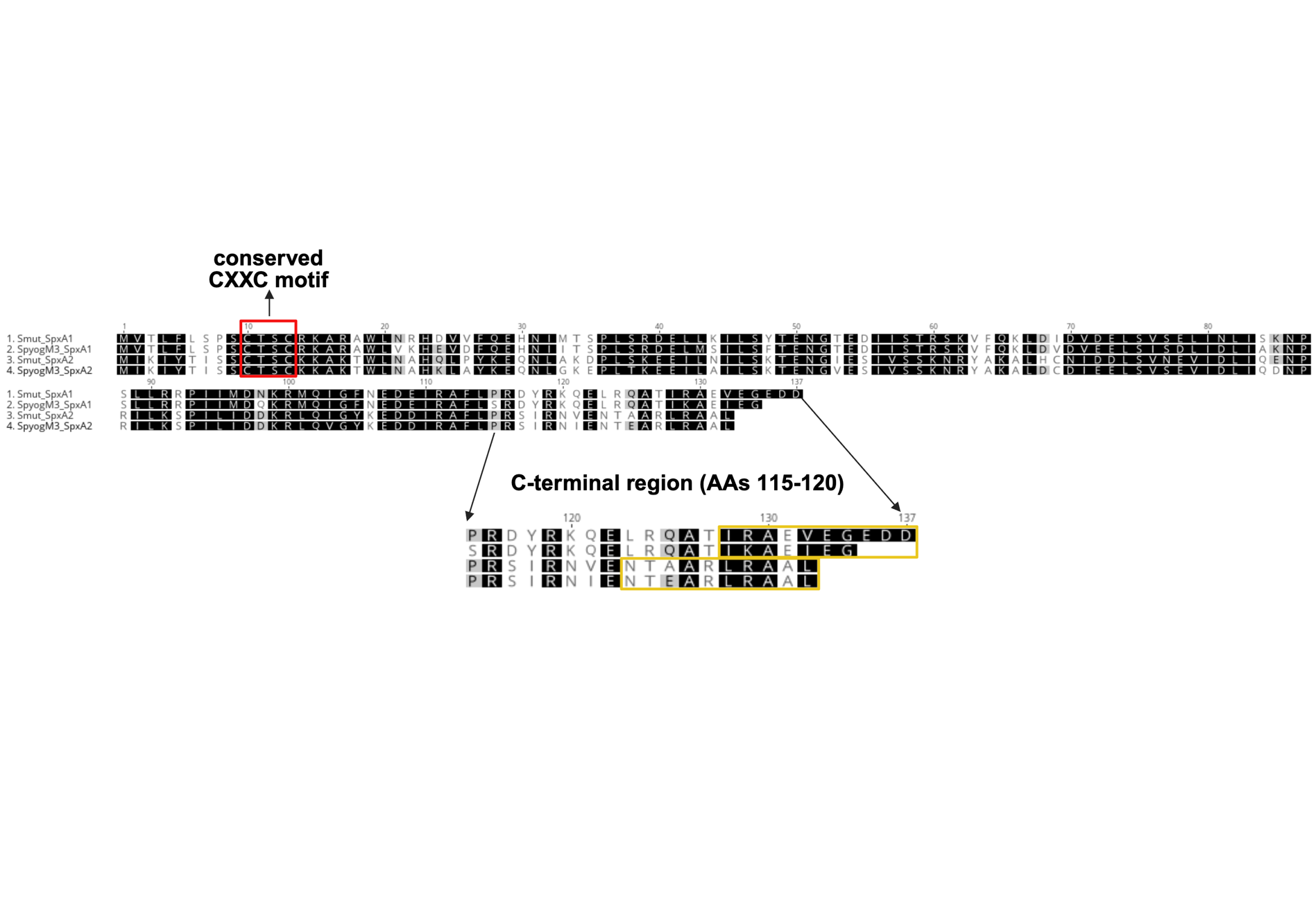

**FIGURE S2**. C-terminal regions of SpxA1 and SpxA2 of GAS show similar amino acid (AA) composition to their *S. mutans* orthologs. Alignments of *S. mutans* and *S. pyogenes* SpxA1 and SpxA2 protein sequences. The conserved CXXC motif between Cys10 and Cys13 is denotated with a red box. The C-terminal region (AAs 115-120) of *S. pyogenes* SpxA1 and SpxA2 (MGAS10870) were “swapped” to create SpxA1 with the SpxA2 C-terminus (A1_A2c) and SpxA2 with the SpxA1 C-terminus (A2_A1c), *see Table ST6 for list of bacterial strains*. The last 10 AAs of *S. mutans* and *S. pyogenes* SpxA1 and SpxA2 are denotated with a yellow box to show AA differences that may contribute to differential degradation rates by the ClpXP protease.

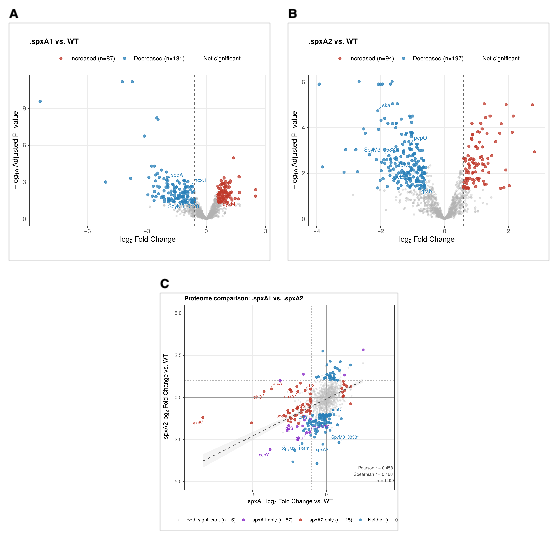

**FIGURE S3**. Global DIA proteomics differential abundance analysis. (**A**) Volcano plot of Δ*spxA1* vs. WT differential protein abundance. Each point represents one detected protein (n=1,199 total). Significantly increased (n=87, blue) and decreased (n=131, red) proteins meet thresholds of *P-*adj < 0.05 and |log_2_FC| ≥ log_2_(1.5) (limma empirical Bayes moderated t-statistic, Benjamini-Hochberg correction). Key proteins discussed in the main text are labeled. (**B**) Volcano plot of ∆spxA2 vs. WT differential protein abundance. Significantly increased (n=94, blue) and decreased (n=197, red) proteins meet the same thresholds as panel A. Key proteins discussed in the main text are labeled. (**C**) Scatter plot comparing log₂ fold-change values across all 1,199 detected proteins in ∆spxA1 vs. WT (x-axis) and ∆spxA2 vs. WT (y-axis). Point color indicates significance category: both significant (black, n=16), ∆spxA1 only (blue, n=57), ∆spxA2 only (pink, n=115), neither significant (gray, n=1,011). Pearson r = 0.458, Spearman r = 0.468, indicating modest positive correlation consistent with shared regulatory contributions alongside substantial paralog-specific effects. Key proteins are labeled. Complete differential abundance results are provided in Supplemental Table ST2.

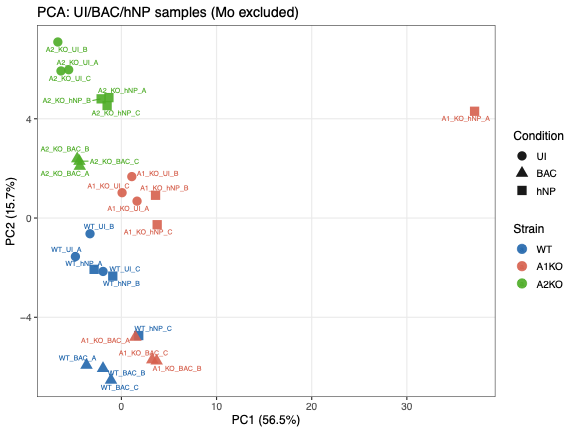

**Figure S4**. NanoString nCounter principal component analysis. Principal component analysis of nSolver-normalized NanoString counts across 27 samples representing WT, Δ*spxA1*, and Δ*spxA2* under uninduced (UI), bacitracin (BAC), and hNP-1 conditions in biological triplicate. PC1 (56.5%) captures the primary variance attributable to stress condition; PC2 (15.7%) reflects strain-dependent differences in transcriptional response. Samples cluster tightly within strain by condition groups confirming high inter-replicate reproducibility. One Δ*spxA1* hNP-1 replicate (A1_KO_hNP_A) was identified as a global expression outlier and excluded from all differential expression analyses (see Supplemental Methods).

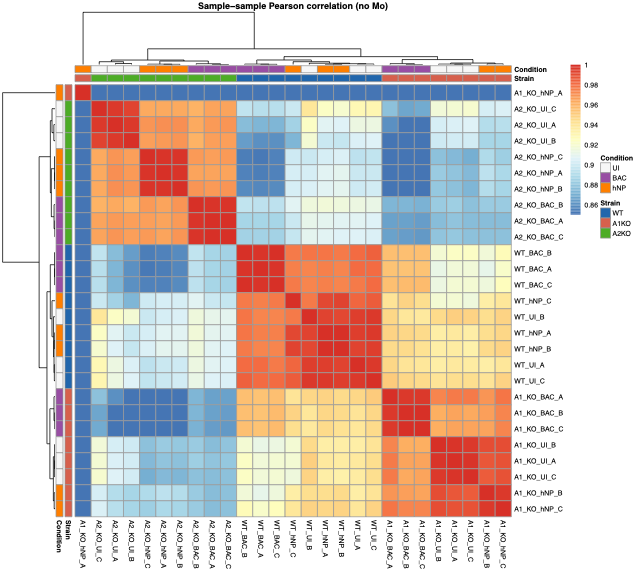

**Figure S5**. NanoString nCounter sample-to-sample Pearson correlation heatmap. Pairwise Pearson correlation of nSolver-normalized NanoString counts across 27 samples representing WT, Δ*spxA1*, and Δ*spxA1* under UI, BAC, and hNP-1 conditions in biological triplicate (macrophage samples excluded). Color intensity reflects pairwise correlation coefficient (range 0.86–1.00); within-group correlations (diagonal blocks) are consistently higher than between-group correlations, confirming strain- and condition-specific transcriptional profiles and high inter-replicate reproducibility. The Δ*spxA1* hNP-1 replicate A (A1_KO_hNP_A) clusters distinctly from the remaining ∆spxA1 hNP-1 replicates, with lower pairwise correlations to all other samples, providing visual confirmation of its identification as a global expression outlier and justifying its exclusion from differential expression analysis (see Supplemental Methods; Supplemental Figure S4).

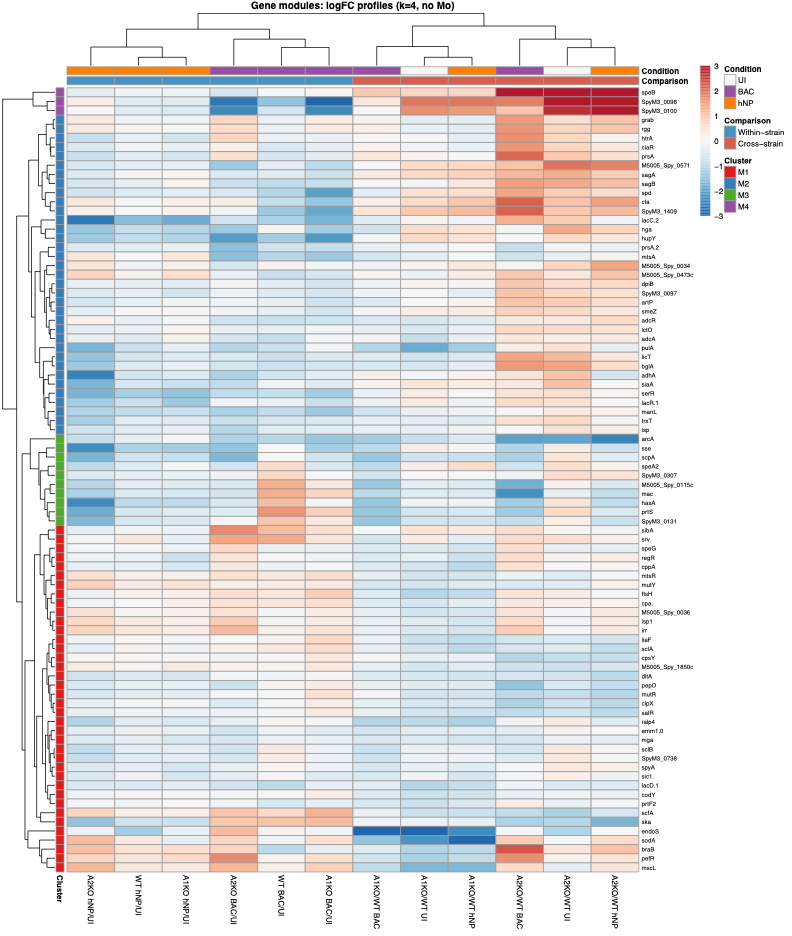

**Figure S6**. NanoString gene module log_2_FC heatmap. Heatmap of log_2_FC profiles across 12 NanoString contrasts for the 86 genes retained for unsupervised hierarchical clustering (|log_2_FC| > log_2_(1.5) in ≥2 contrasts; cross-hybridizing probes excluded). Rows represent individual genes grouped by module assignment (M1, red; M2, blue; M3, green; M4, purple; color sidebar). Columns represent 12 pairwise contrasts grouped by comparison type: within-strain comparisons (BAC/UI and hNP-1/UI for WT, Δ*spxA1*, and Δ*spxA2*) and cross-strain comparisons (Δ*spxA1* vs. WT and Δ*spxA2* vs. WT under UI, BAC, and hNP-1). Color intensity reflects log_2_FC magnitude (scale −3 to +3); red indicates increased expression, blue indicates decreased expression relative to the reference condition. Module assignments were derived from k=4 hierarchical clustering using Euclidean distance with complete linkage; full gene membership and log_2_FC values for all contrasts are provided in Supplemental Table ST5.

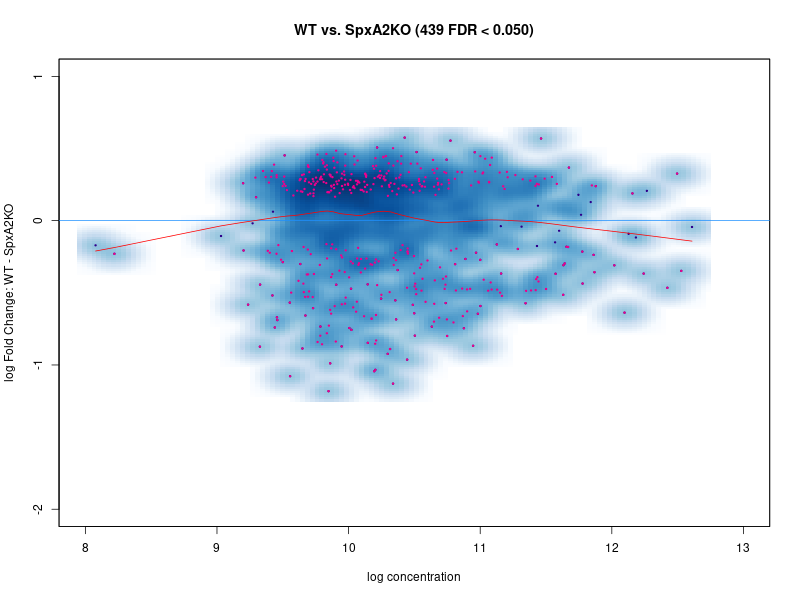

**Figure S7**. ChIP-exo differential CovR binding MA plot – WT vs. ∆*spxA2*. MA plot showing genome-wide differential CovR binding occupancy between MGAS10870 (WT) and Δ*spxA2* from DiffBind analysis. Each point represents one CovR binding site; x-axis shows log concentration (mean normalized ChIP-exo signal across both conditions) and y-axis shows log fold-change (WT − ∆*spxA2*). Magenta points indicate 439 significantly differentially bound sites (FDR < 0.05); black points are non-significant. The red loess curve indicates the overall trend in fold-change across the concentration range. Negative fold-change values indicate higher CovR occupancy in Δ*spxA2* (Δ*spxA2*-gained sites); positive values indicate higher occupancy in WT (MGAS10870-gained sites). DESeq2 normalization was applied within DiffBind v3.20 prior to differential binding analysis (see Supplemental Methods).

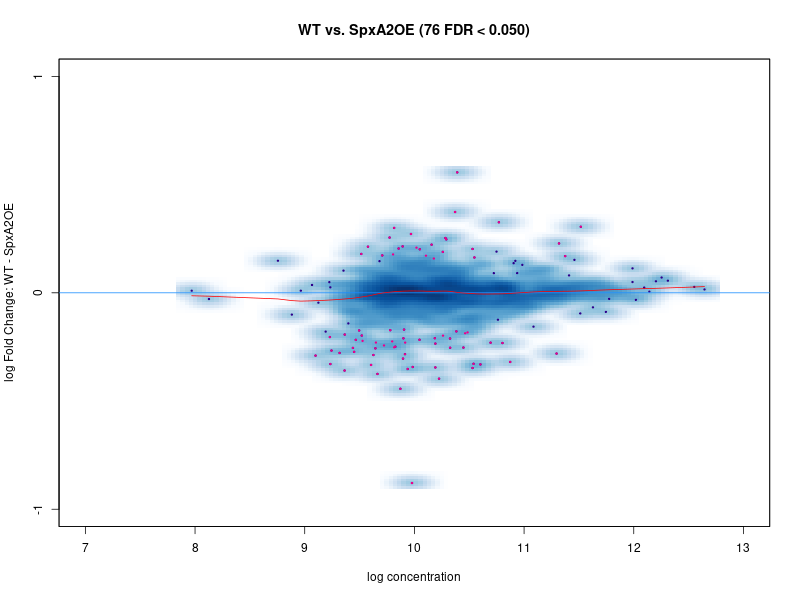

**Figure S8**. ChIP-exo differential CovR binding MA plot – WT vs. LiaS^Q146A^ (SpxA2-OE). MA plot showing genome-wide differential CovR binding occupancy between MGAS10870 (WT) and LiaS^Q146A^ (SpxA2-OE) from DiffBind analysis. Each point represents one CovR binding site; x-axis shows log concentration (mean normalized ChIP-exo signal across both conditions) and y-axis shows log fold-change (WT − SpxA2OE). Magenta points indicate 76 significantly differentially bound sites (FDR < 0.05); black points are non-significant. The red loess curve is near-flat across the concentration range, consistent with the substantially more limited effect of SpxA2 overexpression on CovR occupancy compared to SpxA2 loss (439 differentially bound sites; Supplemental Figure S7). Negative fold-change values indicate higher CovR occupancy in SpxA2-OE; positive values indicate higher occupancy in WT. DESeq2 normalization was applied within DiffBind v3.20 prior to differential binding analysis (see Supplemental Methods).

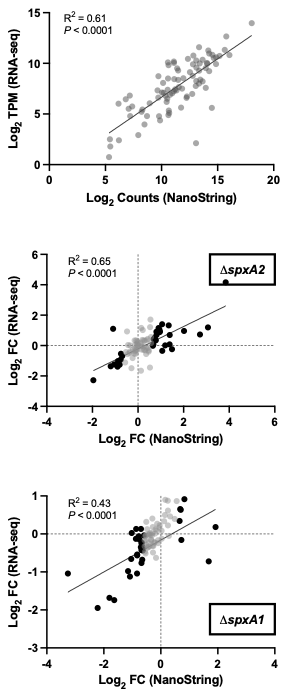

**Figure S9.** Cross-platform validation of NanoString nCounter against RNA-seq. (**A**) Correlation of baseline gene expression levels between NanoString (log_2_ normalized counts, x-axis) and RNA-seq (log_2_ TPM, y-axis) across 86 NanoString panel genes in WT uninduced conditions (R^2^ = 0.61, *P* < 0.0001), confirming concordant quantification of transcript abundance across platforms. (**B**) Correlation of Δ*spxA2* vs. WT log_2_ fold-change values between NanoString (x-axis) and RNA-seq (y-axis) across shared target genes (R^2^ = 0.65, *P* < 0.0001), demonstrating high cross-platform concordance for ∆spxA2-dependent differential expression. (**C**) Correlation of Δ*spxA1* vs. WT log_2_ fold-change values between NanoString (x-axis) and RNA-seq (y-axis) across shared target genes (R^2^ = 0.43, P < 0.0001). The lower R^2^ relative to Δ*spxA2* reflects the smaller Δ*spxA1* regulon and more modest fold-change magnitudes, which are inherently more susceptible to platform-specific variance. The significant positive correlation confirms directional concordance between platforms for Δ*spxA1*-dependent differential expression. RNA-seq and NanoString experiments were performed independently using different biological replicates; concordance reflects cross-platform biological reproducibility.

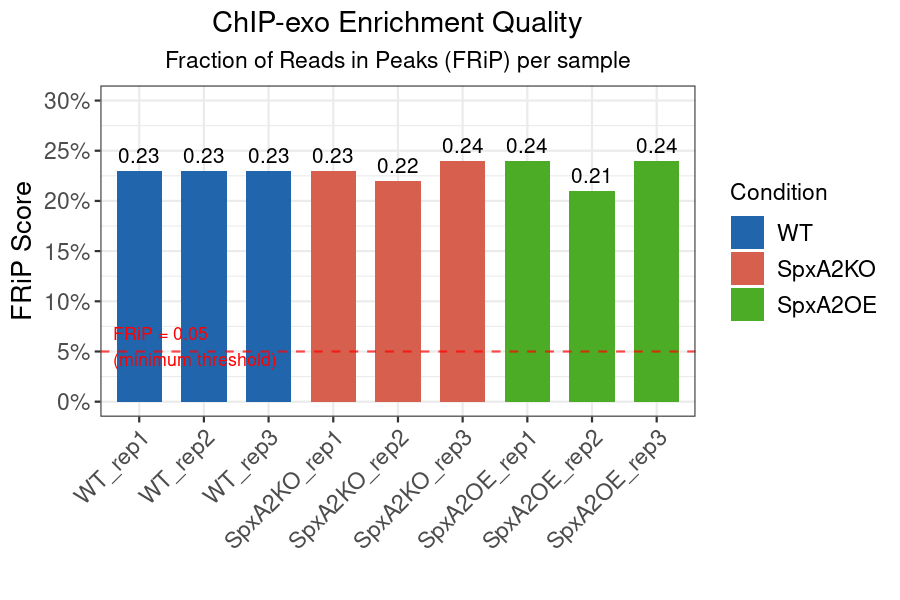

**Figure S10.** ChIP-exo library enrichment quality – Fraction of Reads in Peaks (FRiP) scores. FRiP scores for all nine ChIP-exo libraries across three biological replicates of MGAS10870 (WT, blue), Δ*spxA2* (SpxA2KO, red), and LiaS^Q146A^ (SpxA2OE, green). FRiP scores range from 0.21 to 0.24 across all samples, substantially exceeding the minimum recommended threshold of 0.05 (red dashed line) and confirming high ChIP enrichment quality and efficient λ-exonuclease digestion across all conditions and replicates. The consistency of FRiP scores across strains indicates that differences in CovR binding occupancy detected by differential binding analysis reflect genuine biological differences rather than technical variation in library quality.
